# Distinct roles of nonmuscle myosin II isoforms for establishing tension and elasticity during cell morphodynamics

**DOI:** 10.1101/2020.10.09.333203

**Authors:** Kai Weißenbruch, Justin Grewe, Marc Hippler, Magdalena Fladung, Moritz Tremmel, Kathrin Stricker, Ulrich S. Schwarz, Martin Bastmeyer

## Abstract

Nonmuscle myosin II (NM II) is an integral part of essential cellular processes, including adhesion and migration. Mammalian cells express up to three isoforms termed NM IIA, B, and C. We used U2OS cells to create CRISPR/Cas9-based knockouts of all three isoforms and analyzed the phenotypes on homogeneous substrates, in collagen gels, and on micropatterned substrates. We find that NM IIA is essential to build up cellular tension during initial stages of force generation, while NM IIB is necessary to elastically stabilize NM IIA-generated tension. A scale-bridging mathematical model explains our observations by relating actin fiber stability to the molecular rates of the myosin crossbridge cycle. A dynamic cell stretch/release experiment confirms these predictions and in addition reveals a novel role for NM IIC, namely the ability to establish tensional homeostasis.

## Introduction

The morphodynamics of nonmuscle cells are strongly determined by the contractile actomyosin cytoskeleton, consisting of actin filaments and motor proteins of the nonmuscle myosin II (NM II) class (Burnette et al., 2014; Chen et al., 2010; Gumbiner, 1996; Ingber, 2003; Vicente-Manzanares et al., 2009). The NM II holoenzyme is a hexamer consisting of two heavy chains (NMHC II) that form a homodimer, two regulatory light chains (RLCs), and two essential light chains (ELCs) (Vicente-Manzanares et al., 2009). Phosphorylation of the RLCs mediates the transition from the assembly-incompetent 10S to the assembly-competent 6S conformation of the NM II hexamer (Billington et al., 2013). Once activated, individual NM II hexamers assemble into bipolar filaments of up to 30 hexamers with a typical size of 300 nm, termed myosin minifilaments. These minifilaments can generate tension between antiparallel actin filaments due to their ATP-dependent motor activity. NM II generated forces are then transmitted throughout the cell by subcellular structures such as the actomyosin cortex and different stress fiber subtypes (SFs) (Burnette et al., 2014; Hotulainen and Lappalainen, 2006). Adherent cells are anchored to the extracellular matrix (ECM) at integrin-based focal adhesions (FAs) where high forces can be measured with traction force microscopy (Balaban et al., 2001; Oakes et al., 2017).

Regulation of the actomyosin machinery enables cells to remodel their shape during motion-dependent processes like cell spreading, cell division, or cell migration (Fenix et al., 2016; Svitkina, 2018; Taneja et al., 2020; Vicente-Manzanares et al., 2009; Yamamoto et al., 2019). In general, actomyosin contractility has to be continuously adapted to provide both, short-term dynamic flexibility and long-lasting stability (Ingber, 2003; Mandriota et al., 2019; Matthews et al., 2006). To precisely tune the contractile output, mammalian cells contain up to three different types of myosin II hexamers, which possess different structural and biochemical features. All hexamer-isoforms, which are commonly termed NM IIA, NM IIB and NM IIC, contain the same set of LCs but vary with respect to their heavy chains, which are encoded by three different genes (Heissler and Manstein, 2013). While the cell type-dependent expression, structure, and function of NM IIC is still not clear, the loss of NM IIA and NM IIB causes severe phenotypes in the corresponding KO-mice (Conti et al., 2004; Ma et al., 2010; Takeda et al., 2003; Tullio et al., 1997; Tullio et al., 2001; Uren et al., 2000). In addition, NM IIA and NM IIB are well characterized with respect to their structural and biochemical differences (Barua et al., 2014; Beach et al., 2017; Betapudi et al., 2006; Billington et al., 2013; Sandquist and Means, 2008; Sandquist et al., 2006; Shutova et al., 2012; Shutova et al., 2017; Vicente-Manzanares et al., 2007). NM IIA propels actin filaments 3.5× faster than NM IIB and generates fast contractions (Kovacs et al., 2003; Wang et al., 2000). NM IIB on the other hand can bear more load due to its higher duty ratio (Pato et al., 1996; Wang et al., 2003). Cell culture studies furthermore revealed that NM IIA and NM IIB hexamers co-assemble into heterotypic minifilaments, with a NM IIA to NM IIB gradient from the front to the rear of the cell (Beach et al., 2014; Shutova et al., 2014). Recent publications provide detailed insights that the composition of these heterotypic minifilaments tune contractility during cell polarization and migration (Shutova et al., 2017) or cytokinesis (Taneja et al., 2020). Given such extensive cellular functions and the ubiquitous expression of the NM II isoforms, it is very important to decipher the interplay of the different NM II isoforms for cell shape dynamics and force generation.

Here we address this challenge with a quantitative approach that combines cell experiments and mathematical modelling. Using CRISPR/Cas9 technology, we generated isoform-specific NM II-KO cells from the U2OS cell line, which is a model system for the investigation of SFs (Hotulainen and Lappalainen, 2006; Jiu et al., 2017; Lee et al., 2018; Tojkander et al., 2015; Tojkander et al., 2011). We find that on homogeneously coated substrates, NM IIA-KO cells lack global tension and are unable to build up SFs or mature FAs, while the effect of a NM IIB-KO is less severe and only leads to less well-defined SF networks and smaller FAs. By overexpressing NM IIB in NM IIA-KO cells, we demonstrate that the observed phenotypes do not result from the fact that NM IIA is much stronger expressed than NM IIB. To study the KO cells in a more physiological context, we turned to 3D collagen gels. We find that all KOs can spread within the gels, with the NM IIA-KO cells showing a less contractile phenotype. A gel contraction assay revealed that NM IIA-KO cells are unable to generate global tension while the contraction is only moderately decreased when NM IIB is depleted.

Because these effects are hard to quantify in 3D gels, we next turned to adhesive micropatterns, which are a standard approach for quantitative cell shape analysis in structured environments (Lehnert et al., 2004; Ruprecht et al., 2017; Thery et al., 2006). In previous studies we have investigated the cellular morphology of fibroblasts on dot-shaped micropatterns (Bischofs et al., 2008), where contractile cell types form a contour consisting of a series of concave, inward bent actin arcs (Bischofs et al., 2008; Brand et al., 2017; Kassianidou et al., 2019; Labouesse et al., 2015; Tabdanov et al., 2018; Thery et al., 2006; Zand and Albrecht-Buehler, 1989). Invaginated actin arcs are usually circular and determined by the interplay of surface tension in the cell cortex and line tension in the cell contour, as described mathematically by a Laplace law (Bar-Ziv et al., 1999; Bischofs et al., 2008). A quantitative analysis had revealed a correlation between the spanning distance of the actin arc and the arc radius, suggesting an elastic element in the line tension (tension-elasticity model, TEM). Investigating our NM II-KO cells on cross-shaped micropatterns confirmed the observation from the 3D collagen gels. NM IIA-KO phenotypes are again impaired in tension generation, but adapt their morphology to the structured environment, suggesting that NM IIA is most important for the initial generation of cytoskeletal tension in a homogeneous environment. For NM IIB-KO cells, we find a breakdown of the relation between spanning distance and invaginated arc radius, suggesting that NM IIB is essential to elastically stabilize the NM IIA-generated tension. NM IIC, although expressed in U2OS cells, seems to have hardly no role in cell morphodynamics.

To provide a mechanistic basis for these observations, we developed a mathematical model that bridges the molecular and cellular scales (dynamic tension-elasticity model, dTEM). The main molecular difference between the NM II isoforms are the different rates of their crossbridge cycles. We show that the faster crossbridge cycling of NM IIA leads to a more dynamic generation of tension (higher free velocity, smaller stall force, more dynamic and variable generation of invaginated shapes), thus bridging molecular and cellular scales. To test these predictions, we used our recently published method to measure initial cellular contraction forces and their dynamics during a stretch/release cycle (Hippler et al., 2020). In agreement with our model, we show that NM IIA acts as the initiator of the contractile actomyosin system, while NM IIB is necessary to stabilize these initiated forces to provide long-term stability, especially when cells are under high mechanical stress. In a physiological context, this suggests that the specific properties of both isoforms, NM IIA and NM IIB, are necessary to provide a full and long-lasting force output. Surprisingly, the cell stretch/release experiment also provided a function for NM IIC. Here, the KO cells were not able to perform tensional homeostasis and stay in a contracted mode for prolonged times after stress release.

Together, our results demonstrate that NM IIA, NM IIB and NM IIC have distinct but complementary roles in single cells. While NM IIA is responsible for the dynamic generation of intracellular tension, NM IIB is required to balance the generated forces by adding elastic stability to the actomyosin system. These observations are in excellent agreement with earlier and recent findings that polarized cells form mixed minifilaments with a gradient of NM IIA to NM IIB from front to rear (Beach et al., 2014; Shutova et al., 2014) and that the composition of these heterotypic minifilaments tunes contractility (Shutova et al., 2017; Taneja et al., 2020). In addition, our study revealed a dynamical function of NM IIC, namely a role in establishing tensional homeostasis in a changed mechanical environment.

## Results

### NM II isoforms have a strong effect on stress fiber and focal adhesion formation on homogeneous substrates

To validate the impact of the different NM II isoforms on the cellular phenotype, we used U2OS cells. This cell line serves as a model for the investigation of SFs (Hotulainen and Lappalainen, 2006; Jiu et al., 2017; Lee et al., 2018; Tojkander et al., 2015; Tojkander et al., 2011) and expresses all three NM II isoforms (Figure 1_figure supplement 1A). Since the NMHC II isoforms are encoded by three different genes, it is possible to generate isoform-specific NM II-KO cells. We used CRISPR/Cas9 to target the first coding exons of *MYH9*, *MYH10* and *MYH14,* encoding for NMHC IIA, NMHC IIB and NMHC IIC, respectively. For all NM II isoforms, cell lines lacking the protein of interest were subcultured. The loss of protein expression was confirmed by western blot analysis and immunofluorescence (Figure 1_figure supplement 1A&B). Additionally, DNA sequence analysis revealed frameshifts and pre-mature stop codons in exon 2 of *MYH9* and *MYH10* (Figure 1_figure supplement 1C&D).

We first analyzed the NM II-KO phenotypes on homogeneously coated fibronectin (FN)-substrates and compared our data to previously reported results, where NM II isoforms were depleted via RNAi (Cai et al., 2006; Sandquist et al., 2006; Shutova et al., 2017; Thomas et al., 2015; Vicente-Manzanares et al., 2007) or genetic ablation (Bridgman et al., 2001; Conti et al., 2004; Even-Ram et al., 2007; Lo et al., 2004; Ma et al., 2010; Takeda et al., 2003; Tullio et al., 1997). We visualized SFs and FAs by staining for actin and the FA marker paxillin. Both elements are known to be affected upon interfering with actomyosin contractility (Even-Ram et al., 2007; Sandquist et al., 2006; Shutova et al., 2017; Vicente-Manzanares et al., 2007).

Polarized U2OS WT cells form numerous SFs of different subtypes, as previously described (Hotulainen and Lappalainen, 2006) (Figure 1A). Dorsal SF (dSF) localize along the leading edge and are connected to one FA on the distal end, transverse arcs (tA) align parallel to the leading edge in the lamellum and are not connected to FAs, ventral SF (vSF) localize in the cell center and are connected to FAs on both ends. Depletion of NM IIA leads to a markedly altered cellular phenotype with a branched morphology and several lamellipodia (Figure 1B). No ordered SF-network is built up and only few vSFs remain, dSF or tAs were not observed. The remaining actin structures resemble a dense meshwork of fine, homogeneously distributed actin filaments. In a number of cells, long cell extensions remain, possibly reflecting remnants due to migration defects (Doyle et al., 2012; Even-Ram et al., 2007; Sandquist et al., 2006; Shih and Yamada, 2010). Mature, large FAs are absent in NM IIA-KO cells. Instead, numerous nascent adhesions with a point-like appearance localize along the cell edges, at the remaining vSF, and at the tips of the cell extensions. The effect of the NM IIB-KO is less severe and does not affect overall cell morphology (Figure 1C). All subtypes of SFs are present, but their distinct cellular localization was missing in many cells. In addition, numerous mature FAs were observed throughout the cell body but their localization appeared more diffuse. The depletion of NM IIC did not reveal any phenotypic differences compared to WT cells (Figure 1D).

**Figure 1:**
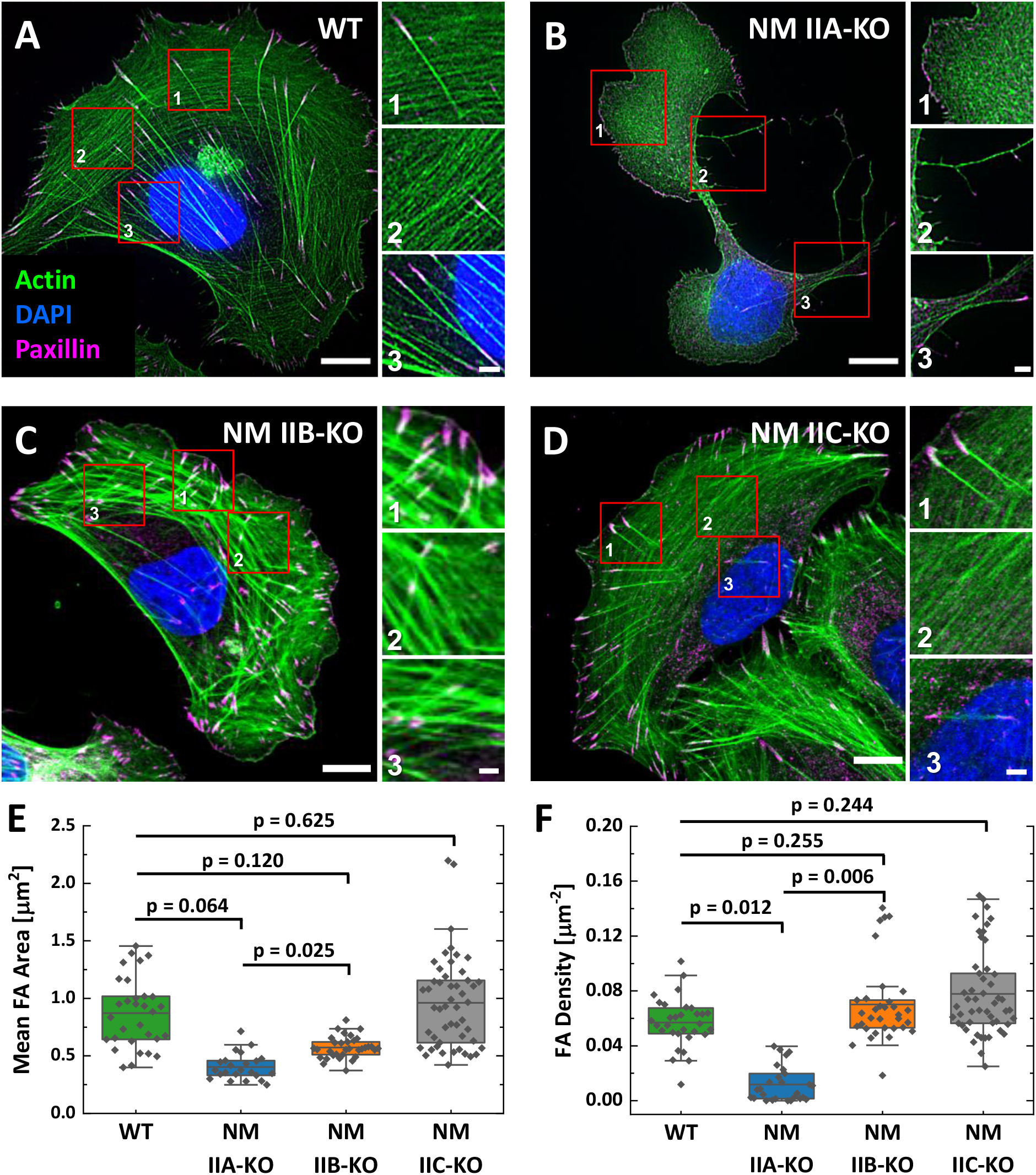
Phenotypes of NM IIA, NM IIB, and NM IIC-KO cell lines are very distinct. **(A)** U2OS WT cells show a polarized phenotype with prominent dorsal stress fibers (dSF) (1), transverse arcs (tA) (2), and ventral stress fibers (vSF) (3). Mature focal adhesions (FA) are visualized by elongated paxillin clusters that localize at the distal ends of dSF or both ends of vSF. **(B)** The NM IIA-KO leads to drastic morphological changes and the loss of most SFs and mature FAs. The overall actin structure resembles a dense meshwork of fine actin filaments (1). At the trailing edge, long cell extensions remain (2) and the only bundled actin fibers resemble vSF (3). **(C)** NM IIB-KO cells reveal slight changes in SF organization and FA structure. dSF (1), tA (2) and vSF (3) are present but their distinct localization pattern is disturbed. **(D)** The phenotype of NM IIC-KO cells is comparable to the WT. dSF (1), tA (2) and vSF (3) localize in a distinct pattern along the cell axis of polarized cells. **(E)** The mean FA area per cell is reduced for NM IIA-KO and NM IIB-KO cells, whereas FA density is only reduced in NM IIA-KO cells **(F)**. Scale bars represent 10 µm for overviews and 2 µm for insets of (A) - (D). **Figure 1_figure supplement 1:** Knockout of NMHC IIA, NMHC IIB and NMHC IIC via CRISPR/Cas9. **Figure 1_figure supplement 2:** FA number and cell area quantification. **Figure 1_figure supplement 3:** Paralog localization in NM II-KO cells. **Figure 1_figure supplement 4:** Intensity quantification of NM II-Paralogs.

We quantified the observed phenotypes by measuring cell area, FA size, and FA density (FA number / cell area). Only FAs ≥ 0.25 µm^2^ were included in the measurements. Cell area does not significantly differ between WT and NM II-KO cell lines, indicating that cell spreading as such is not suppressed by the loss of any NM II isoform (Figure 1_figure supplement 2C). Determination of the mean FA area revealed strongly reduced values for NM IIA-KO cells, a smaller reduction for NM IIB-KO cells, but no reduction for NM IIC-KO cells compared to WT cells (Figure 1E). Concerning FA density, we observed a strong reduction for NM IIA-KO cells, while NM IIB-KO and NM IIC-KO cells do not differ markedly from WT cells (Figure 1F). However, although the total number of FAs was not different, we found that larger FAs occurred less frequently in NM IIB-KO cells, while no difference was observed in NM IIC-KO cells (Figure 1_figure supplement 2A&B).

To investigate if the loss of a certain NM II isoform has an impact on the remaining NM II paralogs, we next compared the localization and intensity of NM IIA-C in WT cells and the respective NM II-KO cell lines (Figure 1_figure supplement 3&4). In polarized WT cells, NM IIA and NM IIC signals are uniformly distributed throughout the cell body whereas NM IIB signals are enriched in the cell center (Figure 1_figure supplement 3A), confirming previous findings (Beach et al., 2014; Kolega, 1998; Shutova et al., 2012; Shutova et al., 2014). Depleting NM IIA strongly alters the localization pattern of both remaining paralogs, NM IIB and NM IIC (Figure 1_figure supplement 3B). Only few NM IIB minifilaments remained and their localization was restricted to the remaining vSFs. The same trend was observed for NM IIC, where a large fraction of NM IIC minifilaments were shifted towards the cell center and clustered along the remaining vSFs. Comparing the intensities of NM IIB or NM IIC minifilaments in NM IIA-KO and WT cells revealed slightly increased but not significantly different values for both paralogs, when NM IIA is depleted (Figure 1_figure supplement 3E and Figure 1_figure supplement 4). In contrast to NM IIA-KO cells, no altered localization of the remaining paralogs was observed in NM IIB-KO cells and only the intensity of NM IIC was slightly but not significantly increased (Figure 1_figure supplement 3C&E and Figure 1_figure supplement 4). No differences were observed for the localization and intensity of NM IIA or NM IIB in NM IIC-KO cells (Figure 1_figure supplement 3D&E and Figure 1_figure supplement 4). Thus, only the loss of NM IIA had an distinct impact on the paralog localization.

Taken together, our results using CRISPR/Cas9-generated U2OS-KO cell lines confirm earlier studies on cell lines derived from KO-mice or on RNAi-mediated knockdown cells (Cai et al., 2006; Even-Ram et al., 2007; Sandquist et al., 2006; Shutova et al., 2017). While the loss of NM IIA leads to drastic morphological changes, the loss of NM IIB only has mild effects. In addition, we could not observe any obvious morphological defects when depleting NM IIC.

### Overexpression of NM IIB does not compensate for the loss of NM IIA

Several studies identified NM IIA as the most abundant expressed isoform, while NM IIB is fairly less strong expressed (Beach et al., 2014; Bekker-Jensen et al., 2017; Betapudi et al., 2011). Given this context, the knockout of NM IIA causes not only the loss of the motors distinct kinetic properties but also a drastic reduction of the totally available NM II hexamers. To determine the ratio of NM IIA to NM IIB minifilaments that assemble in WT cells, we generated fluorescent knock-in cells (Koch et al., 2018), where GFP is expressed under the endogenous promoter of NM IIA or B (Figure 2A). Western blot analysis revealed the heterozygous expression of GFP-NM IIA and B, respectively (Figure 2_figure supplement 1). Therefore, we want to mention that our measurements do not represent absolute numbers of NM IIA or NM IIB molecules but rather give a relative estimation of the ratio of NM IIA and NM IIB hexamers in minifilaments. By measuring GFP signals along segmented actin fibers, we found that the ratio of NM IIA to NM IIB is roughly 4.5:1, proving that NM IIA is indeed the most abundant isoform in minifilaments (Figure 2A&B).

**Figure 2:**
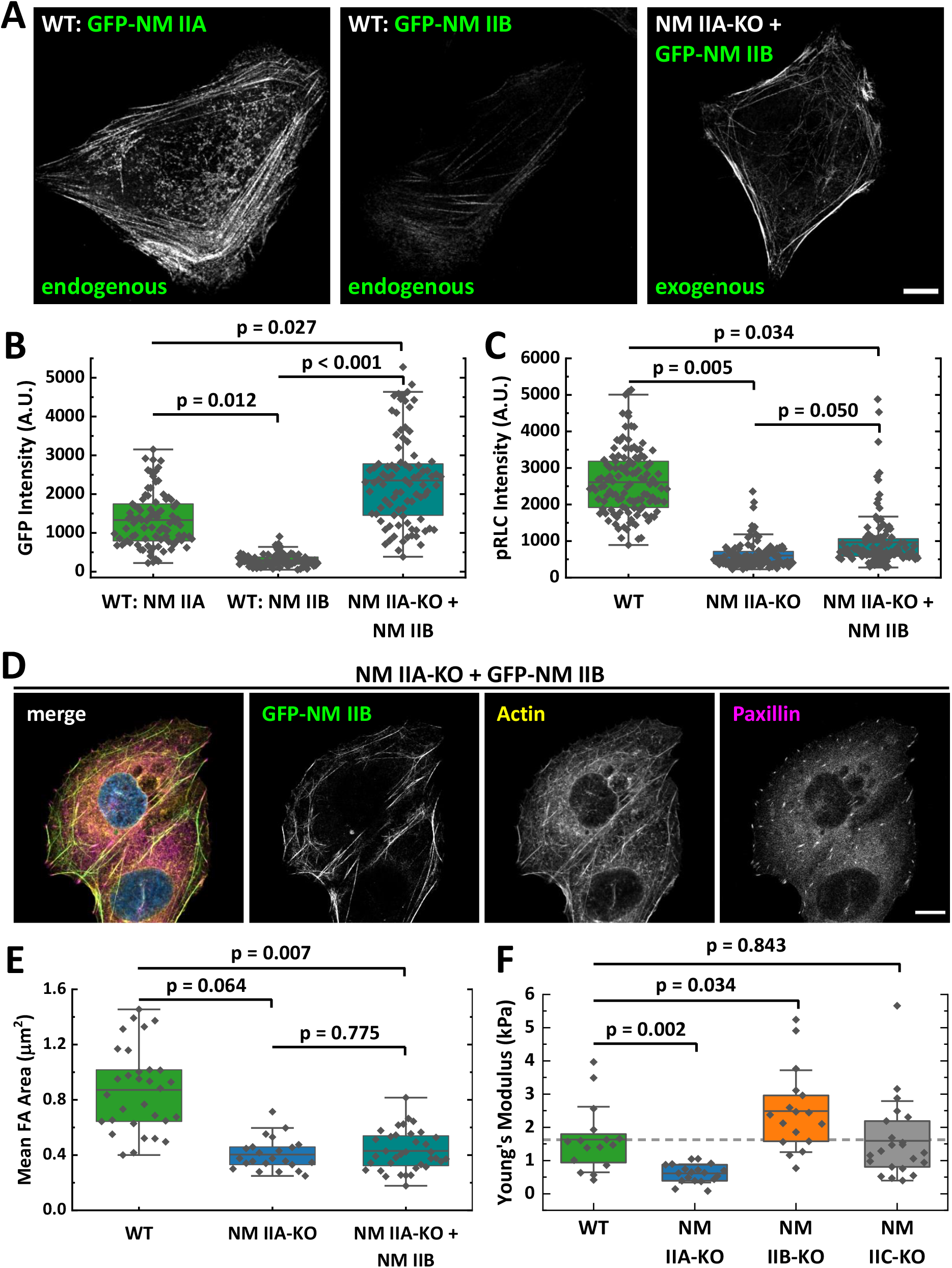
Overexpression of NM II B cannot compensate for the loss of NM II A. **(A)** GFP intensities were measured along segmented actin fibers to compare the ratios of NM IIA and NM IIB in WT and NM IIB overexpressing NM IIA-KO cells. When expressed under the endogenous promoter, the mean GFP intensity of NM IIB is 4.5 times lower compared to NM IIA. To increase the total amount of NM IIB without the interference of NM IIA, NM IIA-KO cells were transiently transfected with exogenous GFP-NM IIB under the CMV promoter. Even in NM IIA-KO cells with high NM IIB expression, NM IIB filament distribution is less homogeneous compared to NM IIA in WT cells. **(B)** Quantitative comparison of endogenous NM IIA and NM IIB in WT cells, and exogenous NM IIB in NM IIA-KO cells. **(C)** pRLC signal intensity was used as a marker for active NM II filaments. The quantitative comparison of WT, NM IIA-KO, and NM IIB overexpressing NM IIA-KO cells shows a substantial reduction of pRLC intensity in NM IIA-KO cells, which is not restored in NM IIB overexpressing NM IIA-KO cells. **(D)** NM IIB overexpression does not phenocopy the WT situation. Immunfluorescent labeling of the actin cytoskeleton (yellow) and the FA marker paxillin (magenta) revealed that SFs are still sparse and FAs are less mature. **(E)** Quantitative comparison of the mean FA size in WT, NM IIA-KO and NM IIB overexpressing NM IIA-KO cells. The values for WT and NM IIA-KO are the same as for Figure 1E and are only shown for comparison. **(F)** AFM nanoindentation experiments were performed to measure the surface tension of the NM II-KO cell lines. Compared to the WT, NM IIA-KO cells possess a significantly lower surface tension, while it is significantly higher for NM IIB-KO cells. No difference was observed for NM IIC-KO cells. Scale bar represents 10 µm in (A) and (D). **Figure 2_figure supplement 1:** Generation of GFP-NM IIA and GFP-NM IIB fusion proteins.

To test whether larger amounts of NM IIB are able to phenocopy the WT situation in NM IIA-KO cells, we overexpressed GFP-NM IIB under a constitutive active promoter and measured again GFP signals along segmented actin fibers. We find that the relative amount of overexpressed GFP-NM IIB is roughly 1.7-fold increased to the amount of endogenous GFP-NM IIA (Figure 2A and B). One should note that these cells also express endogenous NM IIB in addition to the GFP-tagged NM IIB. However, the distribution of NM IIB was still strongly clustered along single actin fibers and does not compare to the distribution of NM IIA in WT cells (Figure 2A). We next used RLCs, phosphorylated at Ser19 (pRLC), as a isoform-independent marker staining for active NM II molecules and measured the fluorescence intensities in WT-, NM IIA-KO-, and NM IIA-KO-cells overexpressing NM IIB-mApple. While the pRLC intensity was drastically reduced in NM IIA-KO cells, NM IIB overexpression only slightly increased the pRLC signal intensities. However, even strongest overexpression of NM IIB in NM IIA-KO cells was not able to fully phenocopy the pRLC level of WT cells (Figure 2C). Similar results were achieved, by comparing the SF formation and FA maturation of GFP-NM IIB overexpressing NM IIA-KO cells to untransfected NM IIA-KO and WT cells (Figure 2D&E). Even high NM IIB levels did not induce the formation of an ordered actin cytoskeleton with bundled SFs of different subtypes. Similarly, FA maturation is still impaired when GFP-NM IIB is overexpressed (Figure 2E). In addition, we performed AFM nanoindentation experiments to compare the surface tension of WT and NM II-KO cells. While NM IIA-KO cells possess a significantly lower surface tension than WT cells, tension is significantly increased in NM IIB-KO cells (Figure 2F). No difference between WT and NM IIC-KO cells was observed. These results are in line with the interpretation that the different kinetic properties of NM IIA and NM IIB tune the contractile properties of the actin cortex and are supported by recently published data obtained by micropipette aspiration assays (Taneja et al., 2020). Thus, not only the absolute quantity of NM II molecules but rather the qualitative properties of both, NM IIA and B, are necessary for the formation of a fully functional actomyosin cytoskeleton.

### Structured environments reveal distinct functions of NM IIA and NM IIB in cellular morphogenesis

To investigate the impact of the NM II-KOs in a more physiological inhomogeneous and structured environment, we next cultured U2OS WT and NM II-KO cell lines in a 3D collagen matrix (Figure 3_figure supplement 1). To probe the contractile properties of our NM II-KO cells, we first cultivated cell seeded collagen gels (CSCG’s) in suspension for 20 hours and measured the gel area at the beginning (red circled area) and the end of the experiment (blue area) (Figure 3_figure supplement 1A). We found that the gel contraction was highest in WT and NM IIC-KO gels, followed by slightly reduced values when using NM IIB-KO cells, and almost no contraction for NM IIA-KO cells. To connect these observations to the cellular morphologies of our NM II-KO cells, we next encased the cells in collagen gels that were attached to the coverslip (Figure 3_figure supplement 1B-E). All cell lines flattened in the collagen gel and in contrast to the cell morphologies on FN-coated coverslips, we now observed phenotypes with concave, inward bent actin arcs that line the cell contour as previously described for various cell types (Bischofs et al., 2008; Brand et al., 2017; Kassianidou et al., 2019; Labouesse et al., 2015; Tabdanov et al., 2018; Thery et al., 2006). This phenotype was most pronounced for WT, NM IIB-KO and NM IIC-KO cells, while the phenotype of the NM IIA-KO cells showed many lamellipodial protrusions. However, these protrusions are also often intersected by small actin arcs.

Since a quantitative evaluation of these actin arcs was not feasible in the structurally ill-defined collagen matrix, we next changed to micropatterned substrates, which allowed us to normalize the cellular phenotypes. We produced cross-shaped FN-micropatterns via microcontact printing, which restrict FA formation to the pattern but still provide a sufficient adhesive area for the spreading of U2OS cells (see methods section for details). Like in collagen gels, all cell lines adapted their shape to the pattern and gained a striking phenotype with concave, inward bent actin arcs that line the cell contour (Figure 3). The actin arcs bridge the passivated substrate areas and have a round shape, which we show to be very close to circular (Figure 3_figure supplement 2). For WT and all three NM II-KOs, circularity errors are very small. The largest deviations were observed for NM IIA-KO cells and the lowest for NM IIB-KO cells. NM II minifilaments localize along the circular actin arcs (Figure 3_figure supplement 3B-D), indicating that they are contractile SFs. From a geometrical point of view, the circularity results from two different NM II-based contributions to cell mechanics: tension in the cortex (surface tension σ) and tension in the actin arcs (line tension λ). Balancing these tensions can explain circular actin arcs with the radius *R* = λ/σ (Laplace law). Typical order of magnitude values have been shown before to be *R* = 10 µm, λ = 20 nN and σ = 2 nN/µm, with the values for λ and σ being extracted from e.g. traction force microscopy on soft elastic substrates or on pillar arrays (Bischofs et al., 2009). In addition, *R* depends on the spanning distance *d* between two adhesion sites, with larger *d* leading to larger *R* values (Bischofs et al., 2008). This dependence can be explained by assuming an elastic line tension λ(*d*) (tension-elasticity model, TEM), suggesting that the mechanics of the peripheral SFs are not only determined by force generating NM II motors, but also by elastic crosslinking, e.g. by the actin crosslinker α-actinin.

**Figure 3:**
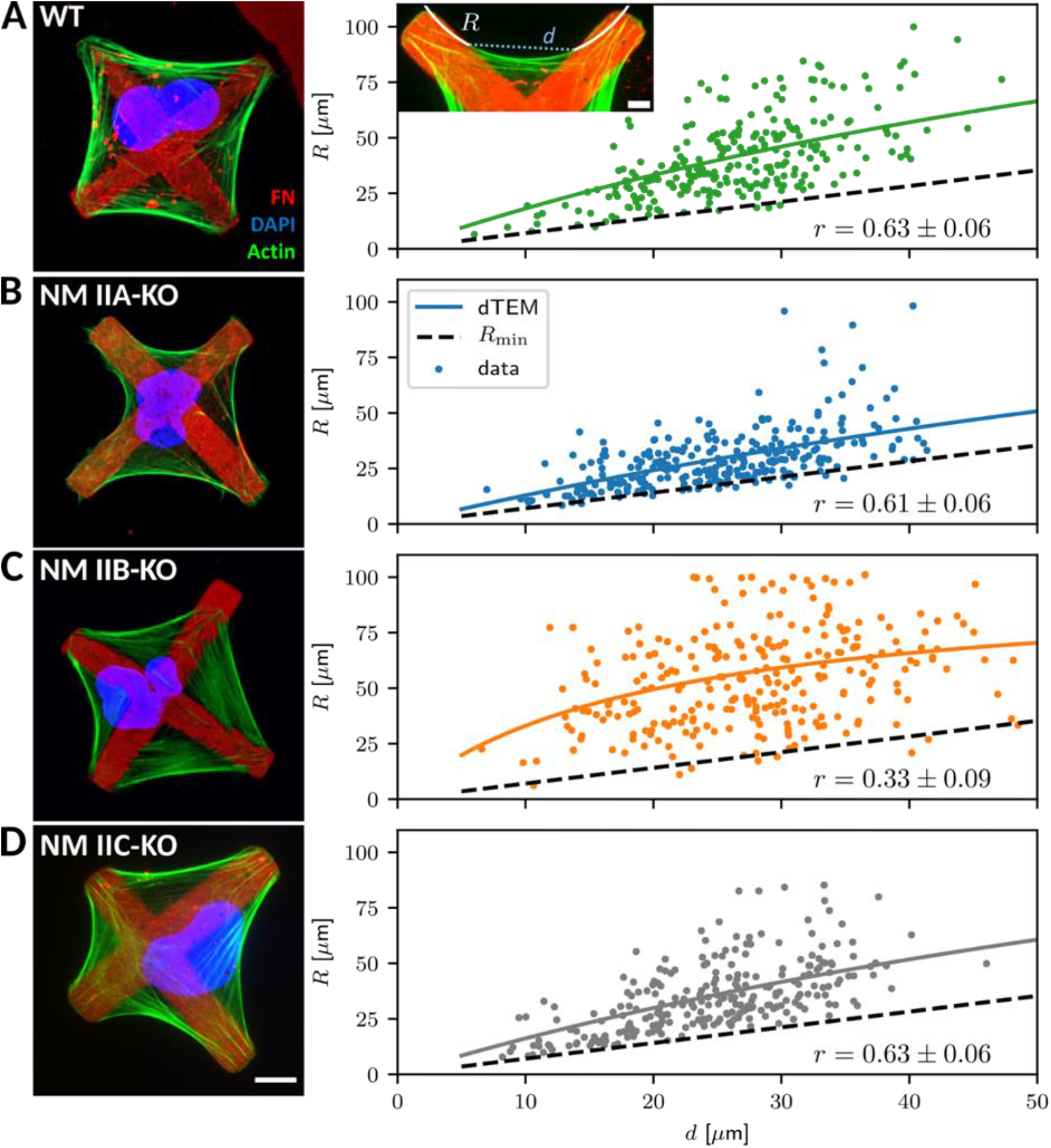
Quantitative shape analysis on cross-shaped micropatterns reveals distinct phenotypes for NM IIA-KO and NM IIB-KO cells. **(A)** U2OS WT cells show prominent invaginated actin arcs along the cell contour, with an invagination radius *R* and a spanning distance *d* (see inset). Quantitative image analysis reveals a positive *R*(*d*) correlation (correlation coefficient r given at bottom right). Solid lines denote the bootstrapped mean fit of the dynamic tension-elasticity model (dTEM), black dashed lines denote the geometrically possible minimal radius. **(B)** U2OS NM IIA-KO cells form invaginated shapes on the cross-patterns despite their strongly perturbed shapes on homogeneous substrates. The spanning distance of the arcs is shorter, but the positive correlation between *R*(*d*) remains. **(C)** Actin arcs of U2OS NM IIB-KO cells are less invaginated compared to WT cells and the *R*(*d*)-correlation is very weak. **(D)** The phenotype of NM IIC-KO cells is comparable to WT and the *R*(*d*) correlation is not affected. Scale bar represents 10 µm for (A) – (D). **Figure 3_figure supplement 1:** NM II-KO phenotypes in collagen gels show prominent actin arcs along the cell contour. **Figure 3_figure supplement 2:** Circularity of invaginated actin arcs. **Figure 3_figure supplement 3**: Co-localization of NM IIA and NM IIB minifilaments along peripheral actin arcs. **Figure 3_figure supplement 4**: NM IIA or B overexpression on micropatterned substrates show the importance of the different motor qualities.

To analyze differences in the NM II-KO cell lines, we measured arc radius *R* and spanning distance *d* and compared their correlation (Figure 3A insert; see methods section for details). WT cells regularly form actin arcs along all cell edges (Figure 3A). Both, NM IIA and NM IIB co-localize with the actin arcs (Figure 3_figure supplement 3B). Quantitative evaluation showed a positive correlation (r = 0.63 ± 0.06) of *R* with increasing *d*, as observed previously (Bischofs et al., 2008; Brand et al., 2017; Tabdanov et al., 2018). Surprisingly, NM IIA-KO cells formed circular arcs and also obeyed a clear *R*(*d*)-correlation (r = 0.61 ± 0.06) (Figure 3B), despite the fact that their phenotype was strongly affected on homogeneously coated FN-substrates. This agrees with our observations in the collagen gels and shows that the structured environment can partially guide the phenotype in the absence of NM IIA-induced contractility. In detail, however, we noticed marked differences compared to WT cells. Although actin arcs along the cell edges are still visible, they do not form as regular as in WT cells. The cell body often covers smaller passivated substrate areas but rather spreads along the crossbars, leading to smaller arcs. Compared to WT cells, fewer internal SFs are present as shown by a weaker image coherency (Figure 3_figure supplement 3A). NM IIB minifilaments co-localize along the actin arcs in NM IIA-KO cells and the pRLC staining is almost completely absent, suggesting that contractile forces are low in these cells (Figure 3_figure supplement 3C). Surprisingly, in NM IIB-KO cells the *R*(*d*) correlation was strongly reduced (r = 0.33 ± 0.09) (Figure 3C), caused by the presence of almost straight arcs that develop independent of the spanning distance *d*. Along these arcs, staining for NM IIA minifilaments and pRLC was comparable to WT cells (Figure 3_figure supplement 3D). We also quantified the degree of internal SF formation but did not find a difference between WT and NM IIB-KO cells (Figure 3_figure supplement 3A). NM IIC-KO cells did not reveal any differences concerning their morphology and the *R*(*d*) correlation was comparable to WT cells (r = 0.63 ± 0.06) (Figure 3D). Importantly, overexpressing GFP-NM IIB in NM IIA-KO cells did not restore the WT phenotype (Figure 3_figure supplement 4A). These cells still spread along the cross bars and do not span over large passivated areas. The *R*(*d*) correlation (r = 0.63 ± 0.06) was comparable to WT and NM IIA-KO cells. Similarly, overexpressing GFP-NM IIA in NM IIB-KO cells did not significantly increase the *R*(*d*) correlation (r = 0.49 ± 0.07) (Figure 3_figure supplement 4B).

In summary, our experimental observations in collagen gels and on cross-shaped micropatterns reveal opposing functions for NM IIA and NM IIB in cell shape determination. NM IIA-KO cells form actin arcs in structured environments, but with small arc radii that are correlated to the spanning distance, while NM IIB-KO cells form actin arcs with large arc radii that are not correlated to the spanning distance.

### NM IIA and NM IIB contribute to dynamic generation of tension and elastic stability, respectively

To better understand these experimental results, we used mathematical models to connect our experimental findings to the molecular differences in the crossbridge cycle with NM IIA generating faster contractions (Kovacs et al., 2003; Wang et al., 2000) and NM IIB bearing more load (Pato et al., 1996; Wang et al., 2003). In contrast to our earlier work, where we developed a static tension-elasticity model (TEM), we now require a dynamical tension-elasticity model (dTEM), connecting the stationary cell shapes to the dynamic crossbridge cycling. We first note that due to geometrical constraints, the circular arcs on our cross-shaped micropattern can have central angles of only up to 90° as shown in Figure 4A, which defines a minimal radius 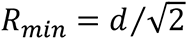 possible for a given spanning distance *d*. We next consider the SF as a dynamic contractile structure that sustains a continuous transport of cytoskeletal material from the FA towards the center of the SF (Figure 4B). This flow can be observed experimentally in vSFs for cells on homogeneously coated substrates and in peripheral arcs for cells on cross-shaped micropatterns (Figure 4_figure supplement 1 and Figure 4_movies 1&2) and, like retrograde flow, is believed to be driven by both actin polymerization in the FAs and myosin-dependent contractile forces (Endlich et al., 2007; Hu et al., 2017; Oakes et al., 2017; Russell et al., 2011; Tojkander et al., 2015). Therefore, it should also depend on the isoform specific motor properties that result from the differences in the crossbridge cycles. Like in muscle cells, mature SFs are organized with sarcomeric arrangements of the myosin motors (Dasbiswas et al., 2018; Hu et al., 2017). Accordingly, the number of serially arranged myosin motors increases linearly with SF length, and SF contraction speed should also increase with length. The stall force *F*_*s*_, however, should not depend on the SF length because in this one-dimensional serial arrangement of motors, each motor feels the same force. A linear scaling between contraction speed and length, as well as the length-independence of the stall force has indeed been observed experimentally in reconstituted SFs (Thoresen et al., 2013). Using an established model for the crossbridge cycle (Figure 4C) and the known differences between the powerstroke rates of NM IIA and NM IIB, we can calculate the stall force *F*_*s*_ for homotypic and heterotypic minifilaments (Grewe and Schwarz, 2020a; Grewe and Schwarz, 2020b). We find that with increasing NM IIB content, the stall force increases and the free velocity decreases, which is mainly an effect of the much smaller duty ratio of NM IIA (Supplemental text and supplement figure S1). For the polymerization at FAs, we assume that its rate increases with force, as has been shown in vitro for mDia1, the main actin polymerization factor in FAs (Jegou et al., 2013). Combining these molecular elements with the geometrical considerations of the TEM (details are given in the supplemental text), we arrive at a surprisingly simple form for the *R*(*d*) relation:

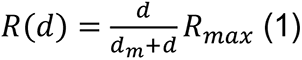

**Figure 4:**
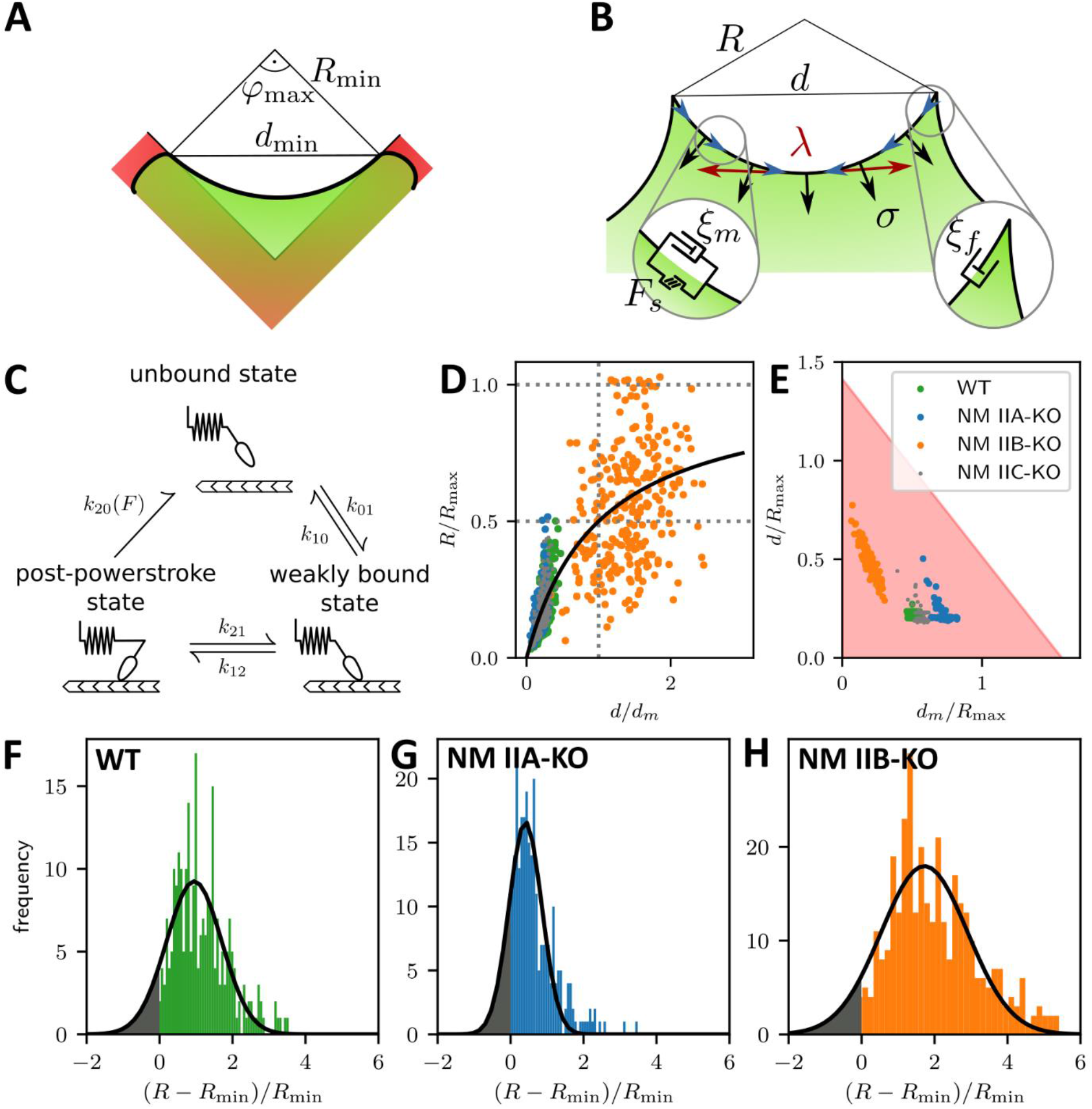
A dynamic tension-elasticity model (dTEM) connects the cellular phenotype to differences in the crossbridge cycling rates. **(A)** Illustration depicting the geometrically possible minimal radius on the cross-shaped micropattern. The circular arcs on our cross pattern can have central angles of only up to 90°. **(B)** Schematics of the mathematical model. At each point along the cell contour, line tension *λ* and surface tension *σ* balance each other and thereby determine the circular arc shape. The insets show the frictional elements required to allow flow of the peripheral fiber (friction coefficients ξ_m_ and ξ_f_ for stress fibers and focal adhesions, respectively). The motor stall force is denoted as F_s_. **(C)** Illustration depicting the three main mechanochemical states during the crossbridge cycle and the corresponding model rates. **(D)** Normalizing experimental results using the fit parameters from Figure 4 yields a master curve. WT, NM IIA-KO and NM IIC-KO cells fall into the linear regime, NM IIB-KO cells into the plateau regime, which corresponds to a loss of correlation. **(E)** *d*_m_/*R*_max_ vs ratio of the maximum of the observed *d*-values to *R*_max_. The region marked in red shows where the central angle of the arc is smaller than 90°. Points denote bootstrapped fit results. **(F-H)** Distributions of differences between observed radius and minimum allowed radius normalized to the minimally allowed radius resemble Gaussian distributions with cut-offs. From this we can estimate the fraction of non-formed rods (grey areas). **Figure 4_figure supplement 1:** Cytoskeletal flow in SFs can be observed on homogeneously coated substrates and cross-shaped micropattern. **Figure 4_movie 1:** Cytoskeletal flow in SFs can be observed on homogeneously coated substrates. **Figure 4_movie 2:** Cytoskeletal flow in SFs can be observed on cross-shaped micropattern.

The maximal radius *R*_*max*_ = *F*_*s*_/*σ* is given by the ratio of stall force *F*_*s*_ and surface tension σ. It can be understood as the arc radius that would be observed if there was no reduction of the tension by the inflow from the FAs and corresponds to the static TEM, with the stall force F_s_ taking the role of the line tension λ in the Laplace law. However, as a NM II-KO is expected to effect both F_s_ and σ to a similar extent, our theory cannot predict directly how R_max_ changes due to the loss of NM II. The spanning distance at half maximal radius d_m_ results from the relative steepness of the force-velocity relations of SFs and FAs (expressed as friction coefficients, compare supplemental text) and defines whether the force is determined by the contraction speed of the fiber or the stall force of the motors. If the spanning distance is small against *d*_*m*_, the observed radius scales linearly with the length of the SF (linear regime), while at spanning distances large against *d*_*m*_, the radius becomes independent of length and is primarily governed by stall force and surface tension (plateau regime).

Fitting eq. (1) to the experimental data shown in Figures 3A-D yield the parameters *R*_*max*_ and *d*_*m*_ for each cell line (solid lines, dashed lines show the minimum radius resulting for a central angle of 90°). The mean fit values and standard deviations for the invaginated arcs are calculated from bootstraps and are listed in table 1. Note that the fit for NM IIB-KO is not entirely meaningful, because a clear correlation is not present in this case. By rescaling the experimental values using the fit parameters, the data points roughly follow a master curve (Figure 4D). For NM IIA-KO, we see that the data is in the linear regime with large d_m_ values, corresponding to the high friction known for NM IIB. This suggests that the measured radii for NM IIA-KO are smaller because the R_max_ cannot be realized given the flow out of the FAs (assuming that λ like F_s_ is increased more than σ for NM IIA-KO, the static TEM would predict the opposite, namely that radii become larger). For NM IIB-KO, Figure 4D shows that the data is closer to the plateau regime (d_m_ small). The small values for d_m_ measured here corresponds to the low friction known for NM IIA. This suggests that the measured radii for NM IIB-KO tend to be larger because the system can in fact dynamically sample R_max_. At the same time, the independence of spanning distance *d* also reflects the breakdown of the correlation, suggesting that NM IIB is required to elastically stabilize the arcs. For NM IIA, we conclude that its main function is to dynamically generate tension, because our data suggest that the small arc radii are mainly determined by its flow properties.

**Table 1:** Mean bootstrapped fit values for the invaginated arcs.

| Cell line | $R_{\max}$ [$\mu\text{m}$] | $d_m$ [$\mu\text{m}$] | $d_m/R_{\max}$ |
| --- | --- | --- | --- |
| WT | $199 \pm 5$ | $100 \pm 5$ | $0.50 \pm 0.02$ |
| NM IIA-KO | $190 \pm 24$ | $137 \pm 22$ | $0.72 \pm 0.04$ |
| NM IIB-KO | $98 \pm 18$ | $20 \pm 9$ | $0.19 \pm 0.05$ |
| NM IIC-KO | $194 \pm 19$ | $110 \pm 14$ | $0.56 \pm 0.03$ |

The error is given as the standard deviation of the bootstrapped fit results. Note that the fit was bounded such that $R_{\max} < 200 \mu\text{m}$ . Therefore, in cases where $R_{\max}$ is close to $200 \mu\text{m}$ , the fit only gives reasonable error estimates for the quotient $d_m/R_{\max}$ .

To further separate the different phenotypes, we plot our data in the two-dimensional parameter space of (*d*_*m*_/*R*_*max*_, *d*/*R*_*max*_) (Figure 4E). The shaded region denotes allowed values due to the central angle being smaller than 90°. Strikingly, the ratio *d*_*m*_/*R*_*max*_, which scales linearly with the ratio of SF friction and motor stall force, increases with the relative amount of NM IIB in the SF, from NM IIB-KO, over WT and NM IIC-KO cells to NM IIA-KO cells. This agrees with our theoretical finding that the ratio of friction and stall force is proportional to the average dwell time of NM II on actin during the crossbridge cycle (Grewe and Schwarz, 2020a; Grewe and Schwarz, 2020b), which is much larger for NM IIB compared to NM IIA. Together these results again suggest that the larger dwell time of NM IIB translates directly into larger radii.

Our results for the NM IIA-KO cells in Figure 4E are closest to the edge of the region with the theoretically permissible arcs, which suggests that in general some arcs cannot form because of geometrical constraints. Our model allows us to further investigate this aspect. Figures 4F-H show that the distribution of the difference of observed radius to the minimum radius, normalized to the minimum radius approximately follows a Gaussian distribution that is, however, cut off at zero difference. Assuming that the missing part of the distribution corresponds to the fraction of arcs that have not formed, we find that there should be approximately 10%, 18% and 7% non-formed arcs for WT, NM IIA-KO and NM IIB-KO cells, respectively. This again suggests that NM IIA is the most important isoform for the formation of arcs, while NM IIB is more important for stabilization.

### NM IIA induced tension and NM IIB derived elastic stability cooperate in the contractile output of cellular stress responses

We next investigated how the loss of NM II isoforms affects cellular contraction forces and the mechanoresponse upon extrinsic stretches (Figure 5). We applied our recently established method for the mechanical stimulation of single cells (Hippler et al., 2020). In brief, 3D micro-scaffolds composed of four non-adhesive walls, each with an inward directed protein-adhesive bar to guide cell attachment (schematically depicted in Figure 5C) were used to measure initial contraction forces. Cells cultured in these scaffolds attach to the bars, thus forming a cross-shaped morphology and pull the walls inwards. These movements were traced with time lapse microscopy for 1 hour, before the cells were detached by trypsinization (Figure 5A). Comparable to traction force microscopy on 2D substrates (Balaban et al., 2001), the displacement values were used to calculate the traction forces exerted by the cells and are in the following given as the sum of all four bars for each cell (Hippler et al., 2020). U2OS WT cells generated a mean initial force of 94 nN (Figure 5B and Figure 5_movie 1), while NM IIA-KO cells generated almost no traction forces (Figure 5_movie 2 and Figure 5_figure supplement 1B) with a mean value of 11 nN (Figure 5B). When measuring the initial forces of NM IIB-KO cells, we obtained a mean value of 112 nN (Figure 5B, Figure 5_movie 3 and Figure 5_figure supplement 1C), which did not significantly differ from WT cells. NM IIC-KO cells also showed no significant difference to the WT (Figure 5_movie 4 and Figure 5_figure supplement 1D) with a mean force value of 110 nN (Figure 5B).

**Figure 5:**
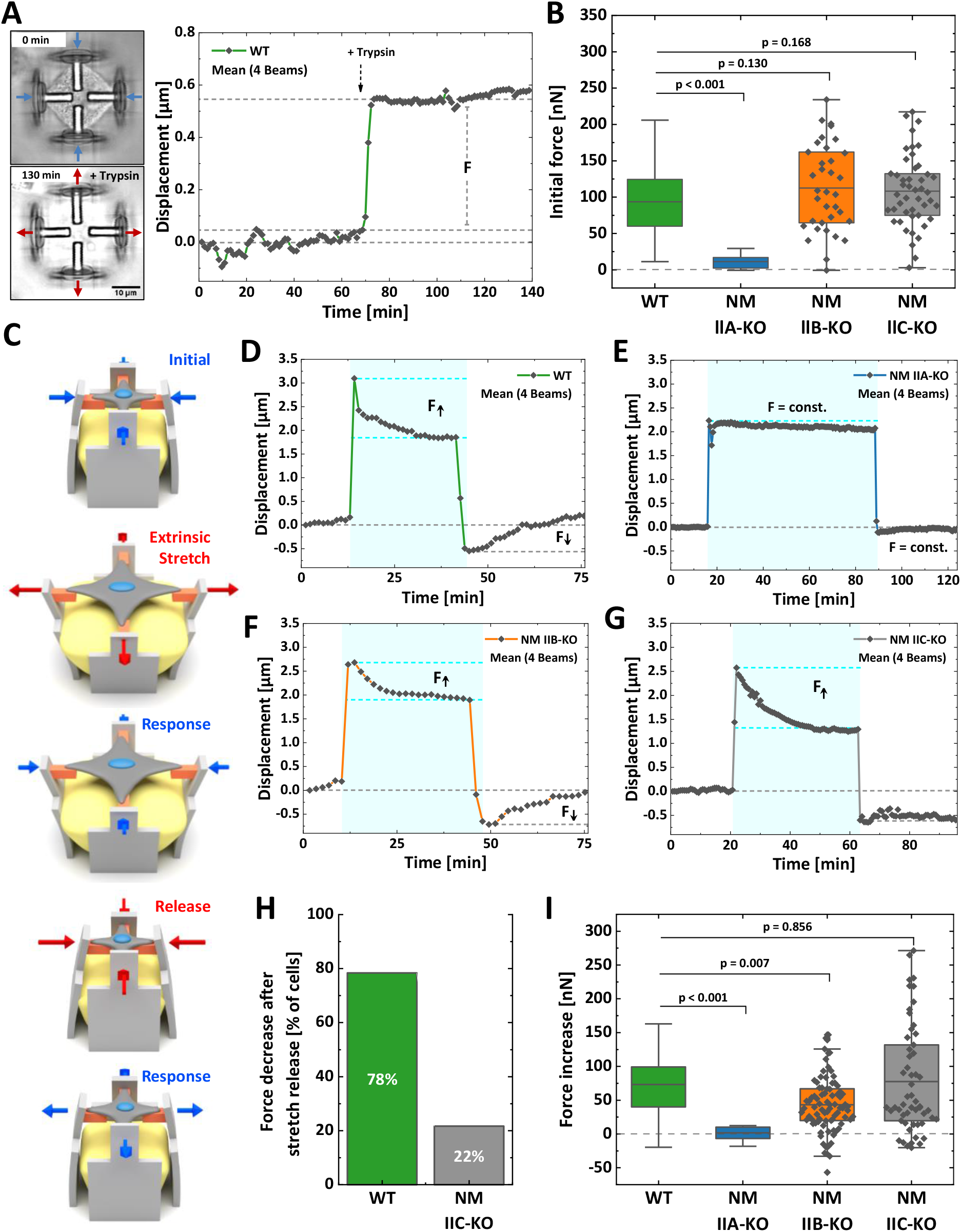
Differential contributions of NM II isoforms to cellular mechanoresponse. Cells were cultured 3D micro-scaffolds composed of four non-adhesive walls, each with an inward directed protein-adhesive bar to guide cell attachment. Cells attach to the bars, form a cross-shaped morphology and pull the walls inwards. **(A)** Initial cellular tractions forces were determined by detaching the cell from the scaffold using trypsin/EDTA and measuring the corresponding average beam displacement as indicated in the plot. **(B)** Comparison of the initial forces of the different cell lines. Data for WT and NM IIA-KO have been published earlier (Hippler et a., 2020), therefore only the mean values are shown. No significant differences were observed between WT (mean value = 94 nN), NM IIB-KO (mean value = 112 nN), and NM IIC-KO cells (mean value = 110 nN). A significant decrease was observed for NM IIA-KO cells (mean value = 11 nN). **(C)** Illustration depicting the stretch-release cycle applied to the cells. **(D-G)** Examples of average beam displacements (corresponding to Figure 5_supplement movies 5-8) are plotted as a function of time. The blue area depicts the time frame, in which the corresponding cell was stretched. **(D)** WT cells actively counteract the stretch and increase their contractile forces until reaching a plateau after ∼ 30-40 min. After releasing the stretch, cellular contraction forces remained high, but decreased to the initial level after 20-30 min. **(E)** No cellular force response is observed, when applying the stretch-release cycle to NM IIA-KO cells, even after longer stretch periods. **(F)** NM IIB-KO cells increase their force after stretching and reach a plateau after 30-40 min. The force increase is lower compared to WT cells. After releasing the stretch, NM IIB-KO cells also reduce their forces until the initial set point is reached. **(G)** NM IIC-KO cells increase their force upon the stretch but do not relax to the initial setpoint within the observed timeframe. **(H)** The quantification shows that a force decrease after the stretch release was observed for 78 % of the WT cells but only for 22 % of the NM IIC-KO cells. **(I)** Comparison of the force increase of WT and NM II-KO cells, after being mechanically stretched. Data for WT and NM IIA-KO have been published earlier (Hippler et a., 2020), therefore only the mean values are shown. A mean increase of 73 nN was observed for WT cells and no force increase for NM IIA-KO cells (mean value = 0.29 nN). Compared to WT cells, NM IIB-KO cells display a significantly lower force increase (mean value = 41 nN), while NM IIC-KO cells show a comparable mean value. However, higher variations in the force response are observed for NM IIC-KO cells. Scale bar represents 10 µm in (A). **Figure 5_figure supplement 1:** Initial force measurements of WT and NM II-KO cells. **Figure 5_figure supplement 2:** Individual beam displacments of stretched WT and NM II-KO cells. **Figure 5_movie 1:** Example of an WT cell during initial force measurement. **Figure 5_movie 2:** Example of an NM IIA-KO cell during initial force measurement. **Figure 5_movie 3:** Example of an NM IIB-KO cell during initial force measurement. **Figure 5_movie 4:** Example of an NM IIC-KO cell during initial force measurement. **Figure 5_movie 5:** Example of an WT cell during reactive force measurement. **Figure 5_movie 6:** Example of an NM IIA-KO cell during reactive force measurement. **Figure 5_movie 7:** Example of an NM IIB-KO cell during reactive force measurement. **Figure 5_movie 8:** Example of an NM IIC-KO cell during reactive force measurement.

To investigate the mechanoresponse of single cells upon extrinsic applied forces, the above described 3D micro-scaffolds were complemented with a block of a guest-host based hydrogel (Hippler et al., 2020). Upon a chemical stimulus this hydrogel expands and pushes the walls apart, thus equibiaxially stretching the cell. Since the process is reversible, removing the stimulus causes a release of extracellular forces and a relaxation of the cell (Figure 5C). We applied the following workflow: WT or MN II KO cells were cultivated for 2 hours in the scaffolds to allow for attachment and equilibration of the cells. Then cells were imaged for 15 min before an equibiaxial stretch of ∼ 20% was applied and the response (displacement in µm) was monitored for 30-60 min. After that time, the stretch was released and the cellular response again monitored for 20-40 min (Figure 5D-G). As previously described (Hippler et al., 2020), WT U2OS cells show a typical response to this stretch-release cycle: Cells counteract the stretch by increasing their traction forces over a time course of ∼ 30 min until they settle on a new plateau value (Figure 5_movie 5 and Figure 5D). After releasing the stretch, WT cells decrease their traction forces again and settle at the initial force value. When applying the stretch-release cycle to NM IIA-KO cells, the cells show no reaction, even after increasing the stretch period to 70 min (Hippler et al., 2020) (Figure 5E and Figure 5_movie 6). In contrast, NM IIB-KO cells revealed a clear response to the stretch-release cycle similar to WT cells (Figure 5F and Figure 5_movie 7). NM IIC-KO cells also increased their traction forces upon stretching and the values were comparable to WT cells. However, 80% of the NM IIC-KO cells did not decrease their intracellular forces after release within the monitored timeframe of 30 min (Figure 5G&H, Figure 5_movie 8 and Figure 5_figure supplement 2D).

As previously described (Hippler et al., 2020), displacements of the scaffolds can be transformed into cellular forces (Figure 5H). Quantifying the force increase (within the blue boxes in Figure 5D-G) showed a mean value of 72 nN for WT cells and no mean force increase for NM IIA-KO cells (Hippler et al., 2020). NM IIB-KO cells also revealed a force increase of 41 nN which was, however, significantly lower than in WT cells (Figure 5H). The force increase of NM IIC-KO cells did not significantly differ from WT cells, but the mean variation was higher. These data strongly support our hypothesis that NM IIA initiates cellular tension, while NM IIB contributes to this tension by providing elastic stability for a stable force output on longer time-scales, i.e. during the mechanoresponse of cells. In addition, NM IIC seems to play a role in the temporal control of the force relaxation, i.e. the mechanical homeostasis of cells.

## Discussion

Here we have systematically analyzed the roles of the three different NM II isoforms for cellular morphodynamics with a combined experimental and theoretical approach. We provide a detailed picture about the complementary functions of NM IIA and NM IIB: generation of dynamic tension by NM IIA and elastic stabilization by NM IIB. U2OS NM II-KO cells cultured on homogeneously coated substrates confirmed previous reports about NM IIA and NM IIB (Even-Ram et al., 2007; Sandquist et al., 2006; Shutova et al., 2017; Vicente-Manzanares et al., 2011; Vicente-Manzanares et al., 2007). Without NM IIA, cells lack global tension, SFs and mature FAs. The NM IIB-KO only leads to a less clear distinction between different types of SF and to smaller FAs. The amount of NM IIA in minifilaments is 4.5 times higher than NM IIB in U2OS cells. Overexpressing NM IIB, however, was not sufficient to compensate for the loss of NM IIA, showing that the induced deficits are due to the loss of the distinct motor properties rather than the overall quantity of the NM II population. A precise quantification of the morphological differences on micropatterned substrates revealed marked changes in the relations between the spanning distance and the radius of invaginated actin arcs. NM IIA-KO cells form small arcs that are correlated to the spanning distance and fail to surpass larger passivated substrate areas. NM IIB-KO cells form large actin arcs, which are poorly correlated with the spanning distance. Similar opposing effects were observed in AFM nanoindentation experiments: While the surface tension was reduced in NM IIA-KO cells, it was increased when NM IIB was depleted. Since the phenotypes on the micropattern arise from the interplay of the two tension regimes σ and λ (Bischofs et al., 2008; Brand et al., 2017), we hypothesize that the different kinetic properties of NM IIA and NM IIB tune the contractile and mechanical properties of both tension regimes in the same manner. Although we cannot distinguish the absolute values of NM IIA or NM IIB that contribute to σ or λ, our results show that the loss of NM IIA reduces both values, line tension in the arcs and surface tension in the cortex, while the loss of NM IIB leads to an increase of both tension regimes.

By connecting the experimental results to our dynamic tension-elasticity model (dTEM), we can explain our results by differences in the molecular crossbridge cycles. NM IIA-KO cells still possess NM IIB-derived elastic stability but lack dynamic tension leading to low intracellular forces. Since the generation of contractile actomyosin bundles is a mechanosensitive process (Tojkander et al., 2015), a polarized actomyosin cytoskeleton is missing in NM IIA-KO cells. Without NM IIA motors, NM IIB is too slow to rearrange the contractile forces in accordance with the fast polymerization of actin filaments. Consequently, the arcs on the crosspattern are smaller and more bent inwards, since the actin polymerization rate overpowers the motor stall force and the surface tension increases the curvature. The only remaining SF resemble vSF, because their turnover is lowest (Kumar et al., 2006; Lee et al., 2018). As FAs are also known to mature in a force-dependent manner (del Rio et al., 2009; Schiller et al., 2013; Vicente-Manzanares et al., 2007), only nascent adhesions remain in the cell periphery and at vSF. NM IIB-KO cells in contrast still possess NM IIA minifilaments, which generate sufficient but unbalanced intracellular forces. The low motor stall force of NM IIA overpowers the actin polymerization rate leading to low curvatures of the actin arcs, which arise independent of the spanning distance. In polarized WT cells, NM IIB is enriched in the central part of the cell and stabilizes vSFs, while higher dynamics arise in NM IIA enriched tAs and indirectly connected dSF (Shutova et al., 2017; Vicente-Manzanares et al., 2008). Since this stabilizing function is missing in NM IIB-KO cells, the fast motor dynamics of NM IIA lead to a high degree of tension, independent of the SF subtype. Although the fast and dynamic motor activity of NM IIA is sufficient to induce the mechanosensitive assembly of all SF subtypes, their distinct localization is disturbed. Likewise, loss of NM IIB does not affect the formation of FAs, however, they do not grow to full size, since the actin templates, in particular the vSFs, are not sufficiently stabilized by the cross-linking properties of NM IIB (Vicente-Manzanares et al., 2011).

No phenotypic change or disturbance in the *R*(*d*) correlation was observed when depleting NM IIC. Thus, our results indicate that NM IIC might be less important for the global generation of contractile forces, at least in our cell line. Structural *in vitro* analysis revealed that NM IIC minifilaments are smaller compared to their paralog counterparts (Billington et al., 2013). This could suggest that NM IIC has a role as a scaffolding protein during the formation of higher ordered NM IIA minifilaments stacks (Fenix et al., 2016), comparable to the role of myosin-18B (Jiu et al., 2019). In line with this, it was reported that NM IIA and NM IIC co-localize throughout the whole cell body in U2OS cells (Beach et al., 2014). Concerning other cell types, a role for NM IIC was shown in regulating the geometry of the epithelial apical junctional-line (Ebrahim et al., 2013) and in the regulation of epithelial microvilli length (Chinowsky et al., 2020). Strikingly, Beach and colleagues showed that the phenotypic switch during EMT (epithelial-mesenchymal transition) and the subsequent invasiveness of murine mammary gland cells goes along with a downregulation of NM IIC and an upregulation of NM IIB (Beach et al., 2011). Thus, NM IIC might be of special interest for the structural organization and integrity of epithelial cell sheets.

We confirmed our results in the context of cellular force generation in live cell experiments. Measuring cellular contraction forces in 3D micro-scaffolds revealed a complete loss of forces for NM IIA-KO cells but no reduction for NM IIB-KO and NM IIC-KO cells. Traction force measurements on 2D substrates showed similar results for NM IIA or NM IIB depleted REF52 cells (Shutova et al., 2017). When measuring the cellular response upon a stretch-release cycle, WT cells showed a behavior that was previously described as mechanical homeostasis (Hippler et al., 2020; Webster et al., 2014; Weng et al., 2016). After the stretch, intracellular forces increased by a factor of two and equilibrated on this new setpoint. After releasing the stretch, cellular contraction forces remained high, but decreased to the initial level after 20-30 min. NM IIA-KO cells did not respond at all to the stretch-release cycle (Hippler et al., 2020).

A clear response was observed when stretching NM IIB-KO cells, however, the force increase did not reach the values of WT cells. After releasing the stretch, cellular contraction forces also decreased to the initial level, however, force decrease was accompanied with oscillations of contractile pulses in about 50% of analyzed traces (Figure 5_movie 7 and red trace in Figure 5_figure supplement 2) as compared to 10% in WT cells. Thus, NM IIB seems to regulate the spatiotemporal response of the actomyosin system by stabilizing NM IIA induced tension. We observed the same trend in collagen gels, where NM IIB-KO cells did contract the gel to a lower amount than WT cells, which is in good agreement with results of others (Meshel et al., 2005). Interestingly, NM IIC-KO cells displayed a delayed response after the stretch was released. In about 80% of the analyzed NM IIC-KO cells, contraction forces did not relax to the initial setpoint within the observed timeframe of 30 min, while this was only the case for 20% of the WT cells. Future experiments might reveal, whether the loss of NM IIC leads to a delay in the cellular mechanoresponse by interfering with the global organization of the NM II contractome. Because the cellular function of NM IIC reported here seems not to be directly related to differences in the powerstroke cycle, future advances in our molecular understanding of this isoform are required to develop a corresponding mathematical model.

Our data can explain how the complementary biochemical features of NM IIA and NM IIB cooperate to build up and maintain the actomyosin cytoskeleton. While NM IIA is responsible for the dynamic generation of intracellular tension, NM IIB is required to balance the generated forces by adding elastic stability to the actomyosin fibers. In this way, the cytoskeletal scaffold provides both, short-term dynamic flexibility and long-lasting stability, allowing cells to dynamically adapt their forces and shapes to varying extracellular geometries and topographies. Considering that tension has to be generated before it can be stabilized, the initiation follows a logical order: The guidance of the entire actomyosin system depends on the presence of NM IIA. This motor can quickly repopulate newly formed protrusions and initiate new contraction sites (Baird et al., 2017). In line with this, other studies proposed a templating function for NM IIA, giving rise to heterotypic minifilaments and dynamizing NM IIB (Fenix et al., 2016; Shutova et al., 2017). As it was shown that NM IIA and NM IIB (and possibly also NM IIC) co-assemble in heterotypic minifilaments (Beach et al., 2014; Shutova et al., 2014), this is probably the reason why the NM IIA-KO leads to differences in the paralog localizations. The phenotype of NM IIA-KO cells in collagen gels and on the micropattern showed, however, similar morphological features than WT cells, suggesting that the main function of NM IIA is guidance of the actomyosin system in the absence of external guidance cues. Once the contraction is initiated, NM IIB co-assembles into the preformed contraction site and stabilizes the tension, as this motor is prone to maintain tension on longer timescales by staying longer bound to the actin cytoskeleton (Sandquist and Means, 2008; Vicente-Manzanares et al., 2008). Thus, the stability of the heterotypic minifilaments is facilitated by the relative composition of NM IIA and NM IIB (Kaufmann and Schwarz, 2020). In a physiological context, such a self-assembling system is able to precisely tune the contractile output of single cells and cell collectives. Therefore, it is not surprising that impairments of this system will lead to massive physiological defects, not only during cell polarization (Heuze et al., 2019), migration (Shutova et al., 2017) or division (Taneja et al., 2020), but also during ECM remodeling (Meshel et al., 2005), as exemplarily shown in our collagen assay. Since different studies showed prominent functions of NM IIB during EMT and invasiveness (Beach et al., 2011; Thomas et al., 2015), or the reinforcement of cell-cell adhesion sites (Heuze et al., 2019), our insights should be transferred to the tissue context, e.g. to explain collective migration effects in development, wound healing or cancer (Scarpa and Mayor, 2016; Shellard and Mayor, 2019; Sunyer et al., 2016; Trepat and Sahai, 2018).

## Materials and Methods

### Cell Culture

U2OS WT cells were obtained from the American Type Culture Collection (Manassas, USA). U2OS NM II-KO cell lines and U2OS GFP-NM IIA or B knock-in cell lines were generated as described in the following sections. All cell lines were tested for mycoplasma infection with negative results. For routine cultivation, cells were passaged every 2-3 days and maintained in DMEM (Pan-Biotech #P04-03590) supplemented with 10% bovine growth serum (HyClone #SH3054.03) at 37°C under a humidified atmosphere containing 5% CO_2_. Cells, plated on FN-coated coverslips or micropatterned substrates, were allowed to spread for 3 hours, cells in 3D micro-scaffolds for 2 hours.

### Generation of NM II-KO and GFP-NM II knock-in cell lines

CRISPR/Cas9 was used to generate knock-out and knock-in cell lines. Guide sequences for the respective protein of interest were determined using the online tool “CHOPCHOP” (https://chopchop.cbu.uib.no/). All used guide sequences are depicted in 5′-to-3′ direction in table 2. Oligos for gRNA construction were obtained from Eurofins genomics (Ebersberg, Germany). NM II-KO cell lines were generated according to the guidelines in (Ran et al., 2013), using the single plasmid system from Feng Zhang’s lab (Addgene #62988). All known splice variants of NMHC IIB and NMHC IIC were targeted by the respective sgRNA. To select for transfected cells, 5 µg mL^-1^ puromycin was added 48 h post transfection to the culture medium and the cells were selected for another 48 h. Single cells were derived by limiting dilution and cell colonies were screened for indels and loss of protein expression.

**Table 2:**
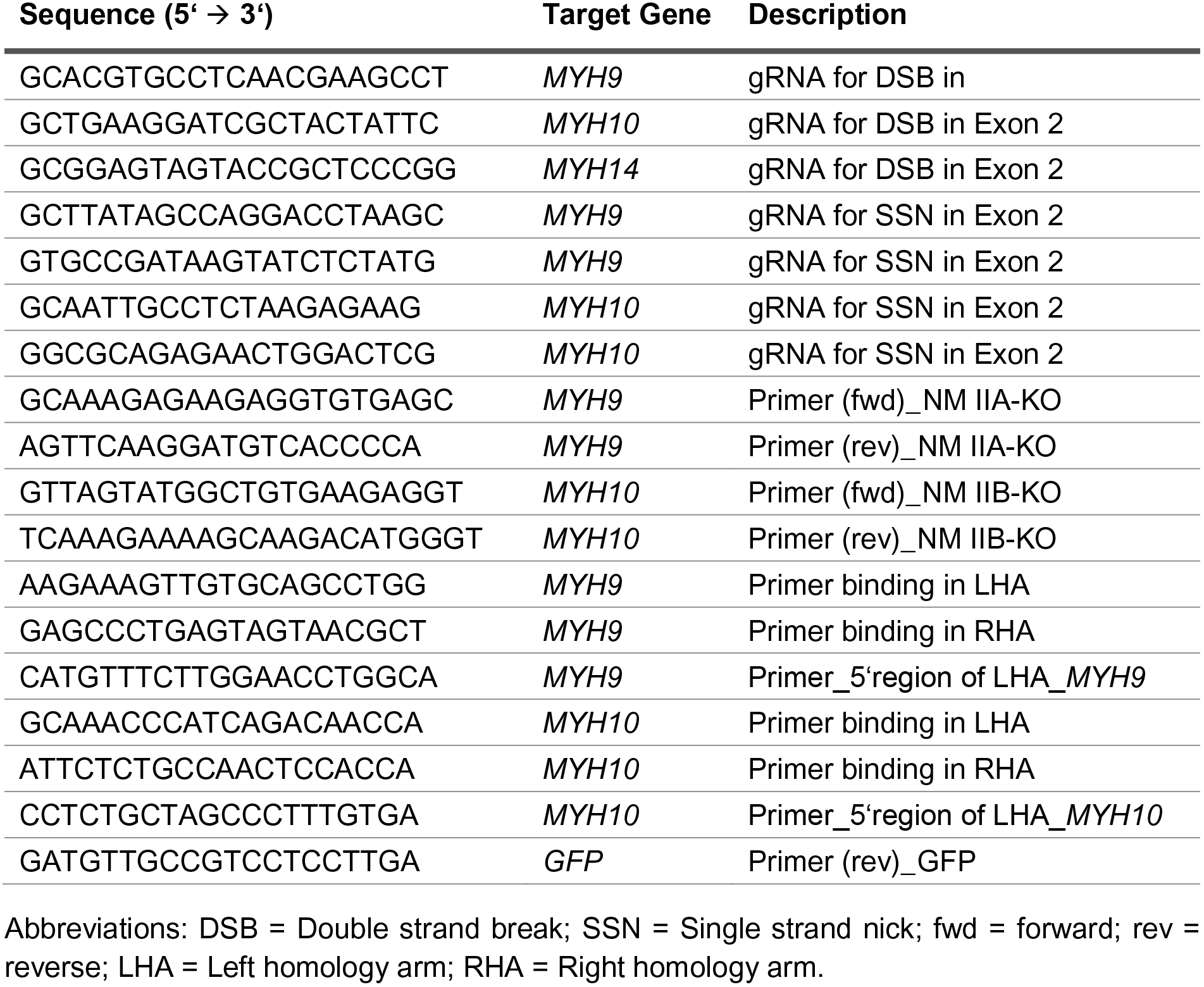
Used gRNA and primer sequences.

Fluorescent knock-in cell lines with GFP fused to the N-terminus of NMHC IIA or NMHC IIB were generated according to the guidelines in (Koch et al., 2018). Briefly, a paired Cas9D10A nickase approach (Ran et al., 2013) was used to generate a double strand break in close proximity of the first coding exon (exon 2) of *MYH9* or *MYH10*. The guide sequences were cloned into pX335-U6-Chimeric_BB-CBh-hSpCas9n(D10A) (Cong et al., 2013). The plasmid was a gift from Feng Zhang (Addgene #42335).

U2OS WT cells were transfected with the according sgRNAs and donor plasmids. GFP-positive cells were sorted using an FACSAria II cell sorter (BD Biosciences). Single cells were derived by limiting dilution and cells were screened for correct insertion of the eGFP by DNA sequence analysis, western blot and immunofluorescence.

### Western blotting

A confluent monolayer of cells in a 6-well plate was lysed in 150 µl ice-cold lysis buffer (187 mM Tris/HCl, 6% SDS, 30% sucrose, 5% β-mercaptoethanol), heated at 95°C for 5 min and centrifuged at 13.000 rpm for 10 min. 30 µl of the supernatant was loaded onto an 8% gel. The proteins were resolved by SDS-PAGE and transferred to a PVDF membrane by tank blotting at 150 mA for 2 h using the Miniprotean III System from Bio-Rad (Hercules, USA). The membrane was blocked for 1 h with 5% skim milk in PBS containing 0.05% Tween-20. The following antibody incubation steps were also carried out in the blocking solution.

Primary antibodies were applied over night at 4°C and secondary antibodies for 2 h at room temperature. Between the antibody incubation steps, membranes were washed in PBS/Tween-20. Following primary antibodies were used: mouse monoclonal to α-Tubulin (Sigma-Aldrich #T5168), rabbit polyclonal to NMHC IIA (BioLegend, #909801), rabbit polyclonal to NMHC IIB (BioLegend, #909901), rabbit monoclonal to NMHC IIC (CST, #8189S), rabbit polyclonal to GFP (Abcam, #ab6556). Secondary horseradish peroxidase-coupled anti-mouse or anti-rabbit antibodies were from Jackson Immunoresearch (#711-036-152 and #715-035-150). The membranes were developed with the SuperSignal^TM^ West Pico PLUS chemiluminescent substrate (ThermoFisher Scientific #34579) according to manufacturer’s instructions. Signal detection was carried out using an Amersham Imager 600 from GE Healthcare (Chicago, USA).

### Sequence analysis

gDNA was isolated using the DNeasy Blood & Tissue Kit (Qiagen #60506) and the target region was amplified via PCR. Primers were designed using the Primer3 freeware tool (Untergasser et al., 2012) and purchased from Eurofins genomics (Ebersberg, Germany). All used primers are listed in table 2. PCR products were either cloned into the pCR II-Blunt-TOPO vector using the Zero Blunt® TOPO® PCR cloning kit (ThermoFisher Scientific #K2875J10) for subsequent sequencing or sequenced directly. Sequencing was carried out at LGC Genomics (Berlin, Germany) and the results were compared to WT sequences using the free available version of SnapGene Viewer (www.snapgene.com/snapgene-viewer).

### Transfection and constructs

Transfections were carried out using Lipofectamine 2000 (ThermoFisher Scientific #11668027) according to manufacturer’s instructions. The cells were transfected 48 h prior to the experiment. CMV-GFP-NMHC IIA (Addgene #11347) and CMV-GFP-NMHC IIB (Addgene #11348) were gifts from Robert Adelstein (Wei and Adelstein, 2000). NMHC IIB-mApple was a gift from Michael Davidson (Addgene #54931). pSPCas9(BB)-2A-Puro (PX459) V2.0 (Addgene #62988) and pX335-U6-Chimeric_BB-CBh-hSpCas9n(D10A) (Addgene #42335) were gifts from Feng Zhang. Guide sequences for the generation of NMHC II depleted cells or GFP knock-in cells were introduced by digesting the plasmids with BbsI and subsequent ligation (Ran et al., 2013). pMK-RQ-*MYH9* and pMK-RQ-*MYH10* donor plasmids for homology directed repair were constructed by flanking the coding sequence of eGFP with 800 bp homology arms upstream and downstream of the double strand break near the start codon of *MYH9* or *MYH10* exon 2 and the plasmids were synthesized by GeneArt (ThermoFisher Scientific).

### Fabrication of micropatterned substrates

Micropatterned substrates were prepared using the microcontact printing technique (Mrksich and Whitesides, 1996). Briefly, a master structure, which serves as a negative mold for the silicon stamp was produced by direct laser writing (Anscombe, 2010) and the stamp was molded from the negative using Sylgard 184 (Dow Corning #105989377). The stamp-pattern resembles a sequence of crosses with different intersections, a bar width of 5 µm and edge length of 45-65 µm. The pattern was either transferred using gold-thiol chemistry (Mrksich et al., 1997) or direct microcontact printing (Fritz and Bastmeyer, 2013). When using gold-thiol chemistry, the stamp was inked with a 1.5 mM solution of octadecylmercaptan (Sigma Aldrich #O1858) in ethanol and pressed onto the gold-coated coverslip, forming a self-assembled monolayer at the protruding parts of the stamp. For the subsequent passivation of uncoated areas, 2.5 mM solution of hexa(ethylene glycol)-terminated alkanethiol (ProChimia Surfaces #TH-001-m11.n6) in ethanol was used. Micropatterned coverslips were functionalized with a solution of 10 µg mL-1 FN from human plasma (Sigma Aldrich #F1056) for 1 h at room temperature. For direct microcontact printing, stamps were incubated for 10 min with a solution of 10 µg ml^-1^ FN and pressed onto uncoated a coverslip. Passivation was carried out using a BSA-Solution of 10 mg ml^-1^ in PBS for backfilling of the coverslip at room temperature for 1 h.

### Fabrication of stimuli-responsive 3D micro-scaffolds

The fabrication and characterization of the stimuli-responsive 3D micro-scaffolds was described in detail in (Hippler et al., 2020). Briefly: A commercial direct laser writing system (Photonic Professional GT, Nanoscribe GmbH) equipped with a 63×, NA = 1.4 oil-immersion objective was used for the fabrication process. In three consecutive writing steps, the various components of the micro-scaffolds were produced by polymerizing liquid photoresists in the voxel of a femto-second pulsed near infrared laser. By using different photoresists that possess hydrophilic or hydrophobic surface properties after polymerization, protein-repellent or protein-adhesive substructures were created. TPETA photoresist was used to write the protein-repellent walls and PETA photoresist for the protein-adhesive beams. For the stimuli-responsive hydrogel, a host-guest based photoresist was polymerized in the center of the scaffold. All mixtures and reagents can be found in (Hippler et al., 2020). Before the sample was used for further experiments, it was immersed overnight in water with 20 mM 1-Adamantanecarboxylic acid (Sigma-Aldrich #106399). This solution triggered the swelling of the hydrogel that helped to remove unpolymerized residues from the material network.

### Immunostaining

Samples were fixed for 10 min using 4% paraformaldehyde in PBS and cells were permeabilized by washing three times for 5 min with PBS containing 0.1% Triton X-100. Following primary antibodies were used: mouse monoclonal to FN (BD Biosciences, #610078), rabbit polyclonal to NMHC IIA (BioLegend, #909801), rabbit polyclonal to NMHC IIB (BioLegend, #909901), rabbit monoclonal to NMHC IIC (CST, #8189S), mouse monoclonal to Paxillin (BD Biosciences, #610619), mouse monoclonal to pRLC at Ser19 (CST, #3675S). All staining incubation steps were carried out in 1% BSA in PBS. Samples were again washed and incubated with fluorescently coupled secondary antibodies and affinity probes. Secondary Alexa Fluor 488-, Alexa Fluor 647- and Cy3-labeled anti-mouse or anti-rabbit antibodies were from Jackson Immunoresearch (West Grove, USA). F-Actin was labeled using Alexa Fluor 488- or Alexa Fluor 647-coupled phalloidin (ThermoFisher Scientific #A12379 and #A22287) and the nucleus was stained with DAPI (Carl Roth #6335.1). Samples were mounted in Mowiol containing 1% N-propyl gallate.

### Fluorescence imaging

Images of immunolabeled samples on cross-patterned substrates were taken on an AxioimagerZ1 microscope (Carl Zeiss, Germany). To obtain high resolution images of minifilaments, the AiryScan Modus of a confocal laser scanning microscope (LSM 800 AiryScan, Carl Zeiss) or a non-serial SR-SIM (Elyra PS.1, Carl Zeiss) were used. The grid for SR-SIM was rotated three times and shifted five times leading to 15 frames raw data of which a final SR-SIM image was calculated with the structured illumination package of ZEN software (Carl Zeiss, Germany). Channels were aligned by using a correction file that was generated by measuring channel misalignment of fluorescent TetraSpeck microspheres (ThermoFischer, #T7280). All images were taken using a 63×, NA = 1.4 oil-immersion objective.

For live cell flow measurements, the incubation chamber was heated to 37°C. Cells were seeded on FN-coated cell culture dishes (MatTek #P35G-1.5-14-C) or micropatterned substrates 3 hours prior to imaging. During imaging, the cells were maintained in phenol red-free DMEM with HEPES and high glucose (ThermoFisher Scientific #21063029), supplemented with 10% bovine growth serum and 1% Pen/Strep.

### Flow analysis and intensity measurements

Flow in vSF or peripheral actin arcs was measured by creating kymographs from a ROI using the reslice function in ImageJ. From these kymographs, movement of individual, persistent minifilaments was tracked manually to determine the flow rate in nm/min.

Quantification of pRLC-, NMHC II-, and GFP-intensities were carried out by calculating the mean intensity along segmented SFs.

### AFM Nanoindentation experiments

We used a NanoWizard AFM (JPK Instruments) equipped with a soft silicon nitride cantilever (MLCT, Bruker) with a nominal spring constant of 0.03 N/m to perform the indentation experiments. The cells were allowed to spread for at least 8h in cell culture dishes (TPP # 93040) before the experiment was performed. For each cell, 16 individual measurements were performed above the nuclear region. A Hertz model was fitted to the resulting force displacement curves and the resulting Young’s moduli were averaged.

### Quantification of FA parameters and *R*(*d*) ratios

Quantification of FAs was performed using the pixel classification functionality of the image analysis suite ilastik (Berg et al., 2019). First, ilastik was trained to mark the cell area. In a separate classification project ilastik was trained to discern between FA and non-FA. The segmentations were exported in the .npy file format for analysis in custom scripts. To determine the number of FAs connected component analysis was applied to the segmented FAs as implemented in openCV 3.4.1.

Quantifications of *R*(*d*) ratios were carried out by manually fitting circles to the peripheral actin arcs of cells on cross-patterned substrates. The spanning distance *d* was defined as the cell area covering the passivated substrate area. In cases, where the cell was polymerizing actin along the functionalized substrate without surpassing the complete distance to the cell edges (as observed in the case of NM IIA-KO cells), only the distance of the cell body covering the passive substrate was considered.

### Stretching experiments and force measurements

Live cell imaging was performed as described above using an LSM 800 equipped with a 40×, NA = 1.2 water-immersion objective and the motorized mechanical stage to sequentially move to all the positions during the time series. To exchange solutions during the experiment, the sample was mounted in a self-built fluidic chamber. For initial force measurements, cells were imaged for 30-60 min under steady-state conditions and then detached from the substrate using Trypsin/EDTA. For stretching experiments, the cells were first imaged for 10 min under steady-state conditions. To induce the mechanical stretch, the solution was exchanged to medium containing 20 mM 1-Adamantanecarboxylic acid and the cells were imaged in the stretched state for 30-70 min. Releasing the stretch was obtained by again replacing the medium with normal imaging medium and the cellular reaction was monitored for up to 30-50 min.

To calculate the initial and reactive forces, the images were analyzed by digital image cross-correlation based on a custom-written MATLAB code (MathWorks) as described in (Hippler et al., 2020). In every scene, regions of interest were defined on the four beams and every frame of the time series was compared to a reference image at t = 0. The calculation of the maximum cross-correlation function resulted in the 2D local displacement vector. Four different positions per beam were tracked and averaged to obtain a mean displacement per beam as a function of time. Additional tracking of solid marker structures and reference scaffolds without cells was used to correct potential offsets and deviations that are not induced by cellular forces. Ultimately, the measured displacements were converted to cell forces by modeling the properties of the micro-scaffolds by finite element calculations (see (Hippler et al., 2020) for details).

### Cell-seeded collagen gels

CSCGs were generated according to the guidelines in (Provenzano et al., 2010). We used collagen I from rat tails (Enzo Life Sciences #ALX-522-435) and seeded 1.5×10^5^ cells in collagen matrices with a final concentration of 1 mg/ml. As a neutralizing buffer, 0.1 M HEPES in 2×PBS was used in equal volumes to the collagen solution. 250 µl total volume were distributed in 18 mm glass bottom dishes and allowed to polymerize at 37°C. After 2h, the dish was backfilled with DMEM and the CSCG’s were cultivated in suspension for another 18h. After 20h total incubation time, CSCG’s were fixed in 4% PFA and the diameter was measured.

For gels fixed to the glass bottom, dishes were pre-coated with thin plate coating collagen I (Enzo Life Sciences #ALX-522-440-0050) and allowed to dry overnight. On the following day, CSCG’s were fabricated as described and polymerized on the pre-coated culture dishes.

### Modeling

For more information about the dTEM and the parameters, we refer the reader to the supplemental text, where a detailed description can be found.

## Acknowledgments

We thank Alisha Rapp (KIT) for her help with the analysis of GFP- and pRLC intensities. This work is supported by the Deutsche Forschungsgemeinschaft (DFG, German Research Foundation) under Germany’s Excellence Strategy through EXC 2082/1-390761711 (the Karlsruhe-Heidelberg 3DMM2O Excellence Cluster, to USS and MB) and EXC 2181/1 - 390900948 (the Heidelberg STRUCTURES Excellence Cluster, to USS). USS is a member of the Interdisciplinary Center for Scientific Computing (IWR) at Heidelberg. JG acknowledges support by the Research Training Group of the Landesstiftung Baden-Württemberg on Mathematical Modeling for the Quantitative Biosciences.

## Figure supplement

**Figure 1_figure supplement 1:**
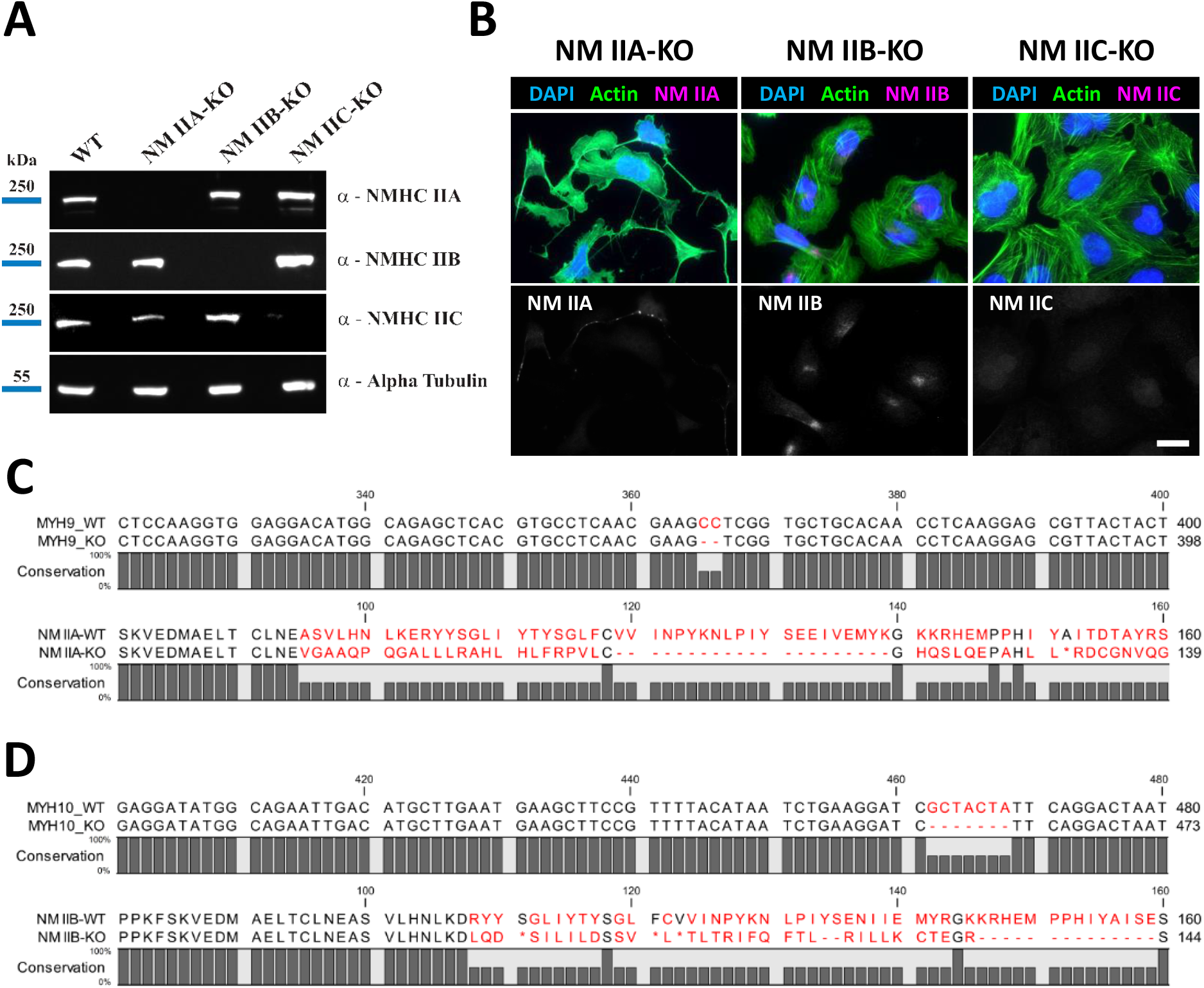
Knockout of NMHC IIA, NMHC IIB and NMHC IIC via CRISPR/Cas9. NM II-KO cell lines were generated via CRISPR/Cas9 and the loss of protein expression was confirmed via western blot **(A)** and immunofluorescent labeling **(B)**. Deletions in the coding sequences of the first coding exons (exon 2) of NMHC IIA **(C)** and NMHC IIB **(D)** lead to frame shifts and premature stop codons in corresponding protein sequences (marked by asteriscs). Scale bar represents 20 µm.

**Figure 1_figure supplement 2:**
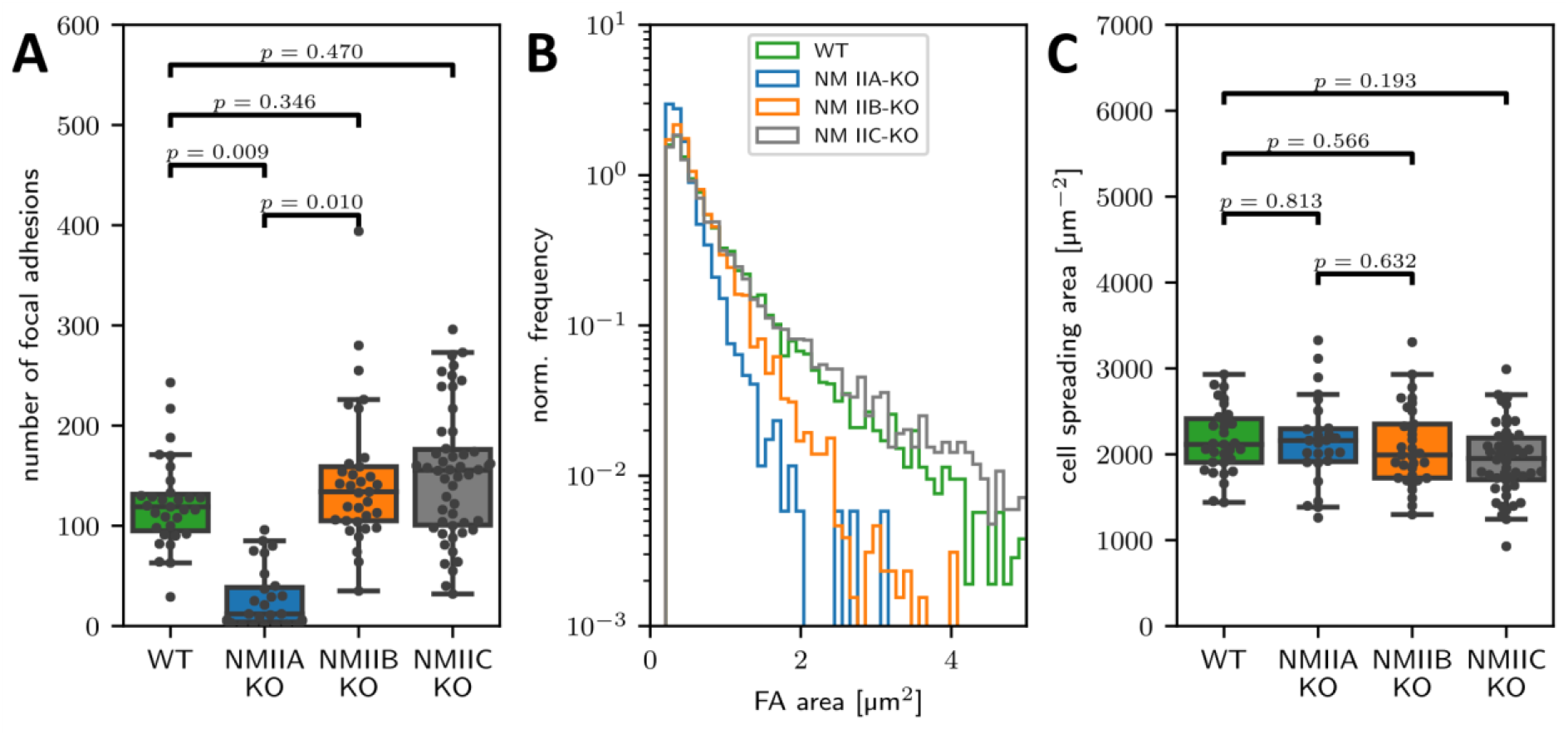
Quantification of FA number and cell spreading area. Quantifications showing the number of focal adhesions **(A)**, their size frequency **(B)**, and the cell area **(C)** of WT and the respective NM II-KO cell lines. **(A)** NM IIA-KO cells show a reduced number of mature focal adhesions (larger than 0.25 µm²), while no difference was observed for NM IIB-KO and NM IIC-KO cells. **(B)** The frequency distribution of FAs is shifted to smaller sizes in NM IIA-KO and NM IIB-KO cells as compared to WT cells. No difference was observed for NM IIC-KO cells. **(C)** The cell spreading area is not affected by the loss of NM IIA, NM IIB or NM IIC.

**Figure 1_figure supplement 3:**
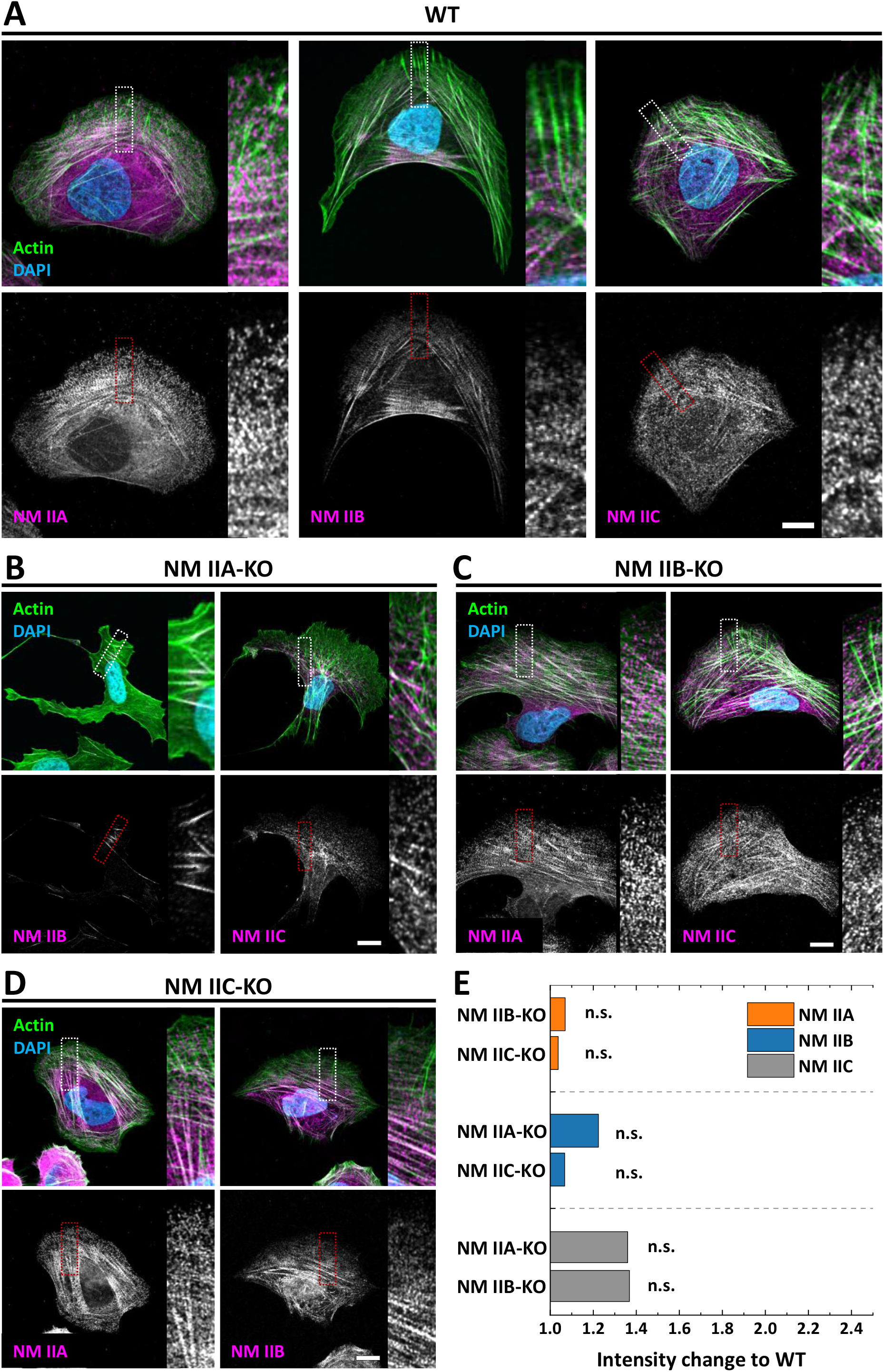
Paralog localization in NM II-KO cells. Immunfluorescent labeling was used to analyze the localization and intensity of the NM II paralogs in U2OS WT or the respective NM II-KO cells. **(A)** In polarized WT cells, all isoforms co-localize with actin fibers throughout the cell body (NM IIA and NM IIC) or in the cell center (NM IIB). **(B)** In NM IIA-KO cells, the remaining paralogs show an altered localization pattern: NM IIB and NM IIC minifilaments are clustered along the remaining SFs in the cell center. **(C)** In NM IIB-KO cells, NM IIA and NM IIC minifilaments still localize throughout the cell body, comparable to WT cells. **(D)** The loss of NM IIC did not change the localization of the remaining paralogs. NM IIA is homogeneously distrubuted and NM IIB accumulates in the cell center. **(E)** The mean fluorescence intensities of NM IIA-C was measured along segmented actin fibers and the fold change to the WT was calculated. All scenarios resulted in only mild intensity increases and no significant differences were observed. Scale bars represent 10 µm in (A)-(D).

**Figure 1_figure supplement 4:**
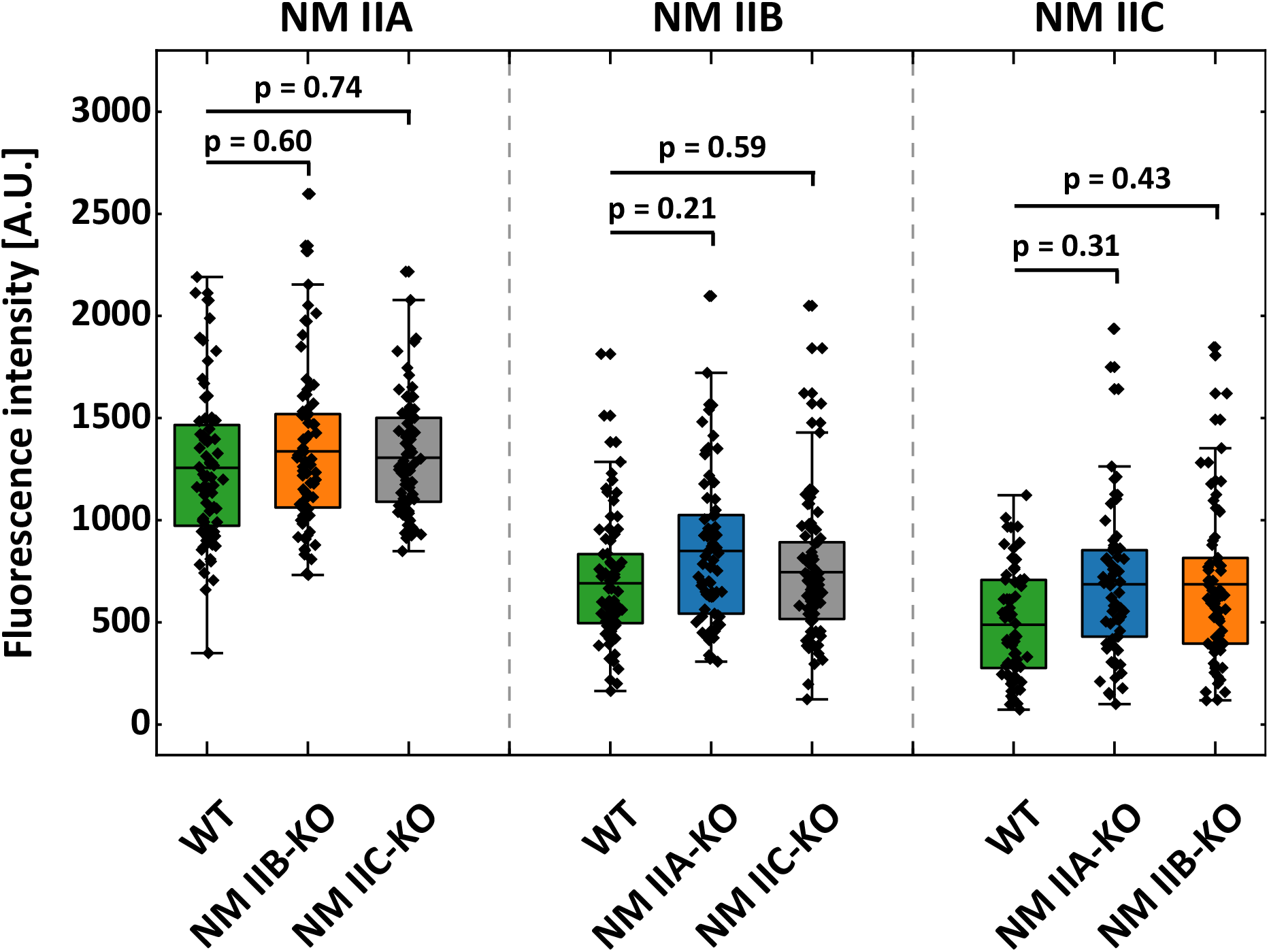
Intensity quantification of NM II-Paralogs. Segmented actin fibers were used to determine the fluorescence intensities of NM IIA-C in WT and the respective NM II-KO cells. Shown are the absolute intensity values, used to calculate the fold changes in Figure 1_figure supplement 3E.

**Figure 2_figure supplement 1:**
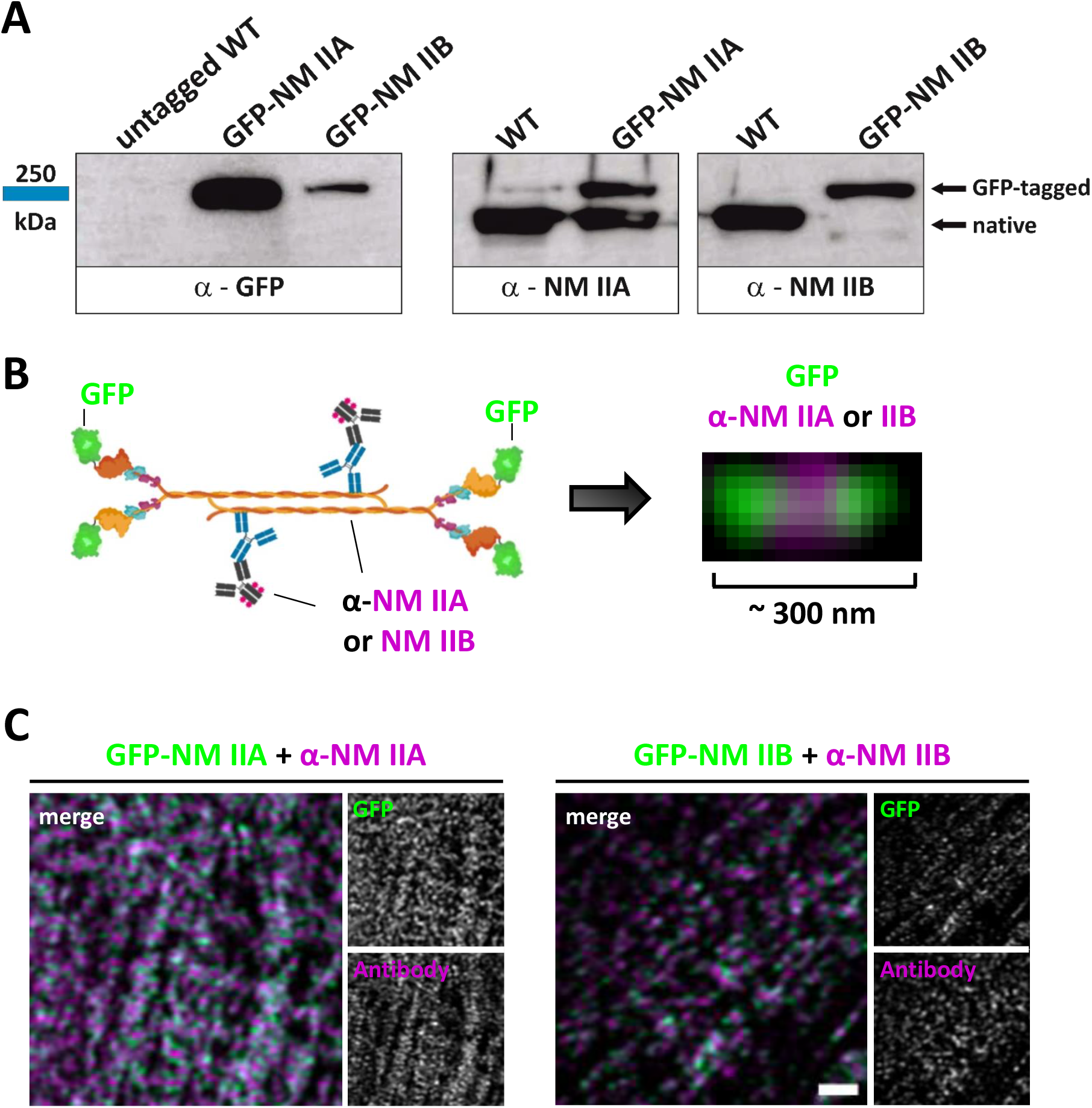
Generation of GFP-NM IIA and GFP-NM IIB fusion proteins. GFP-NM IIA or GFP-NM IIB reporter cell lines were generated according to the guidelines in (Koch et al., 2018). **(A)** Western Blots depicting the insertion of the GFP tag in the knock-in cell lines and heterozygous expression of GFP with NM IIA or NM IIB. **(B)** Illustration depicting localization of the GFP tag and antibody binding sites in bipolar minifilaments. The GFP sequence was introduced at the 5’-End of the coding sequences, resulting in N-terminally tagged NM IIA or NM IIB fusion proteins. The antibodies for NM IIA or NM IIB bind at the C-terminal region, resulting in the depicted bipolar staining pattern. **(C)** Exemplary images of bipolar minifilaments in GFP-NM IIA or GFP-NM IIB cells. Scale bar in (C) represents 1 µm.

**Figure 3_figure supplement 1:**
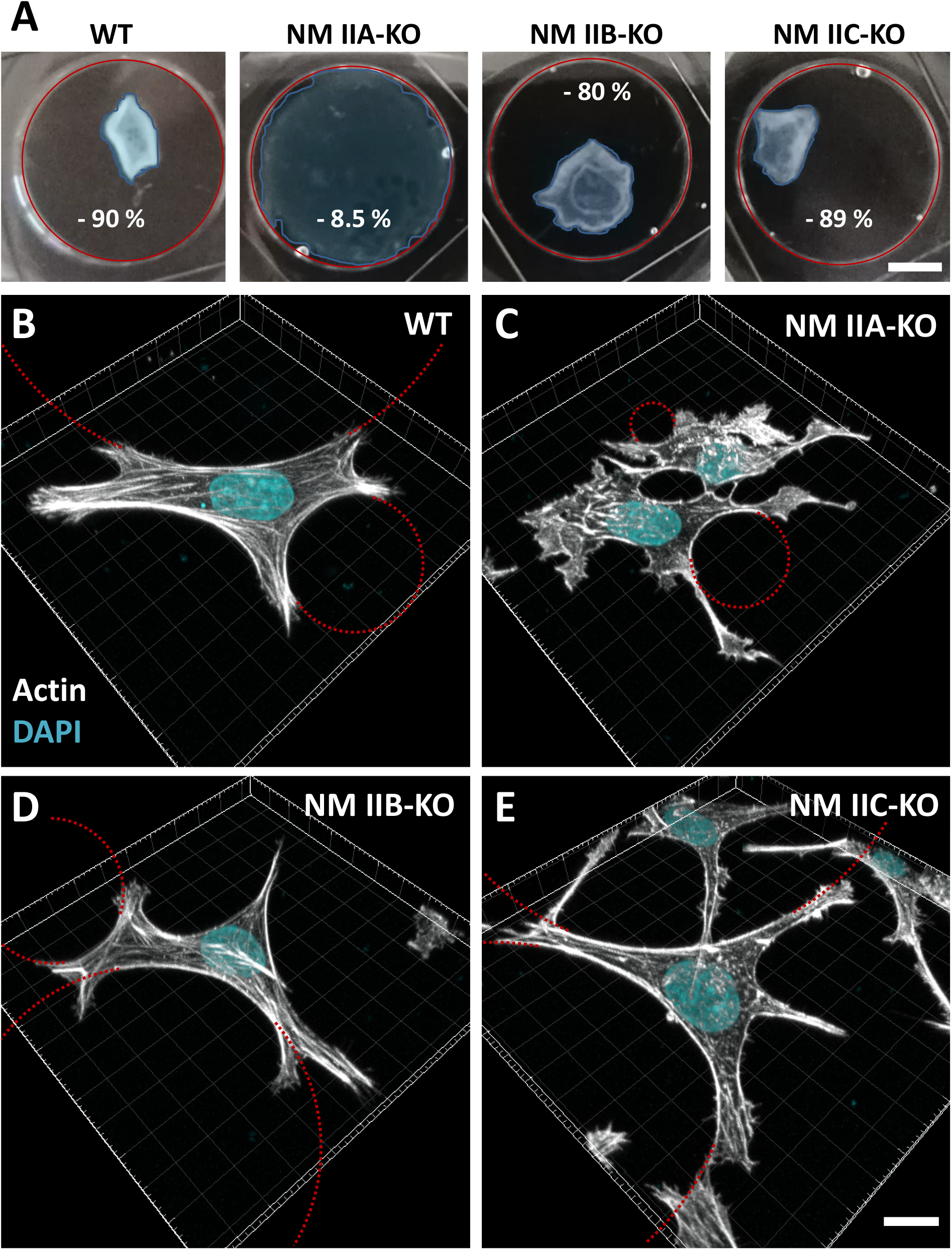
NM II-KO phenotypes in collagen gels show distinct contractile behaviors with prominent actin arcs along the cell contour. **(A)** Cell seeded collagen gels (CSCG’s) were cultivated in suspension and the area was measured at the beginning and the end of the experiment. Red circles indicate the initial area, the blue area shows the CSCG after a total incubation time of 20 hours. WT cells contract the CSCG to 10% of the initial area. NM IIA-KO cells barely contract the gel and 91.5% of the initial size is still present after the incubation time. For NM IIB-KO cells, the contraction was also lower than in the WT case with 20% of the initial area left. No difference was observed for WT and NM IIC-KO cells. **(B-E)** 3D images of WT and NM II-KO cells, encased in collagen gels that were attached to the coverslip. All cell lines flattened in the collagen meshwork and showed prominent actin arcs along parts of the cell contour. For the sake of clarity, only some of the actin arcs were marked by the fitted circles in red. While WT, NM IIB-KO and NM IIC-KO show comparable phenotypic features, the phenotype of NM IIA-KO cells is characterized by many lamellipodial protrusions and extensions of various shape and size. However, these protrusions are also often intersected by small actin arcs. Scale bars represent 5 mm for (A) and 15 µm for (B – E).

**Figure 3_figure supplement 2:**
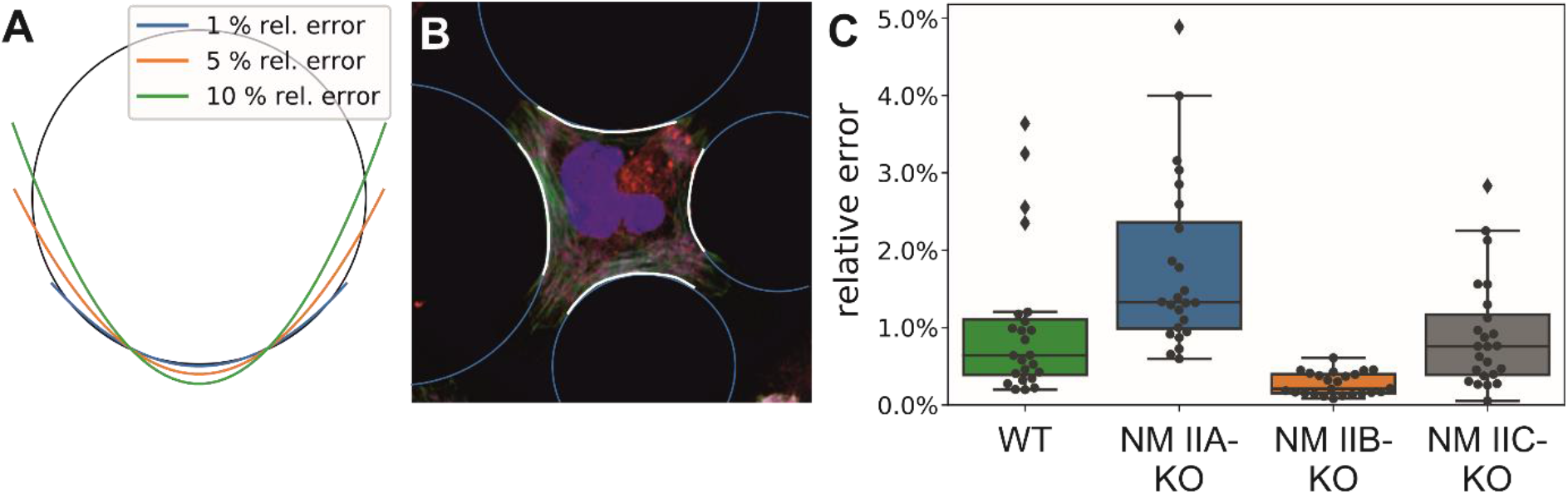
Circularity of invaginated actin arcs. **(A)** The root mean squared deviation of the distance from points on the arc to the fitted center relative to the fitted radius is a measure for the circularity of the arc, as demonstrated here for parabolic curves.**(B)** Cell contours were fitted with the ImageJ/Fiji-plugin JFilament (Smith et al., 2010) and then fit to circular arcs with a custom-written software using the hyper least squares algorithm (Kanatani and Rangarajan, 2011). **(C)** For WT and all three KOs, circularity errors are relatively small, with typical values around 2% and few outliers above. The largest deviations are observed for NM IIA-KO.

**Figure 3_figure supplement 3:**
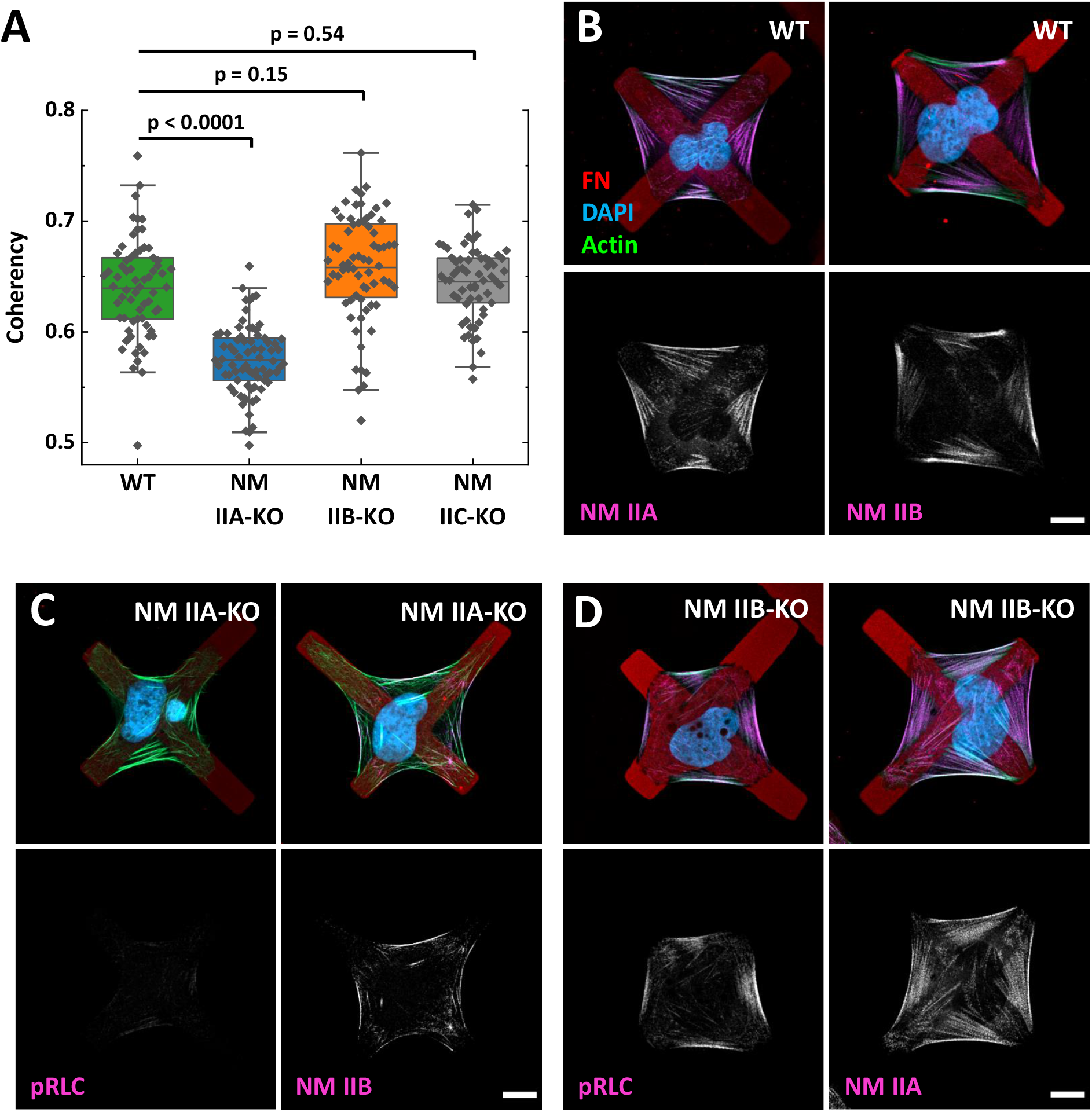
Localization of NM IIA and NM IIB minifilaments along peripheral actin arcs. **(A)** The amount of actin fibers is quantified by the coherency of the structure tensor measured for the cell interior with a custom-made python script. Only NM IIA-KO cells show a significantly weaker coherency, demonstrating that these cells don’t develop internal SFs. **(B)** In WT cells, NM IIA and NM IIB minifilaments localize along peripheral actin arcs and internal SFs. **(C)** In NM IIA-KO cells, the pRLC signal is almost absent and NM IIB minifilaments appear mainly along the actin arcs. **(D)** In NM IIB-KO cells, NM IIA minifilaments are homogeneously distributed along the actin arcs and the internal SFs. The pRLC staining is comparable to the WT cells. Scale bars represent 10 µm for (A)-(C).

**Figure 3_figure supplement 4:**
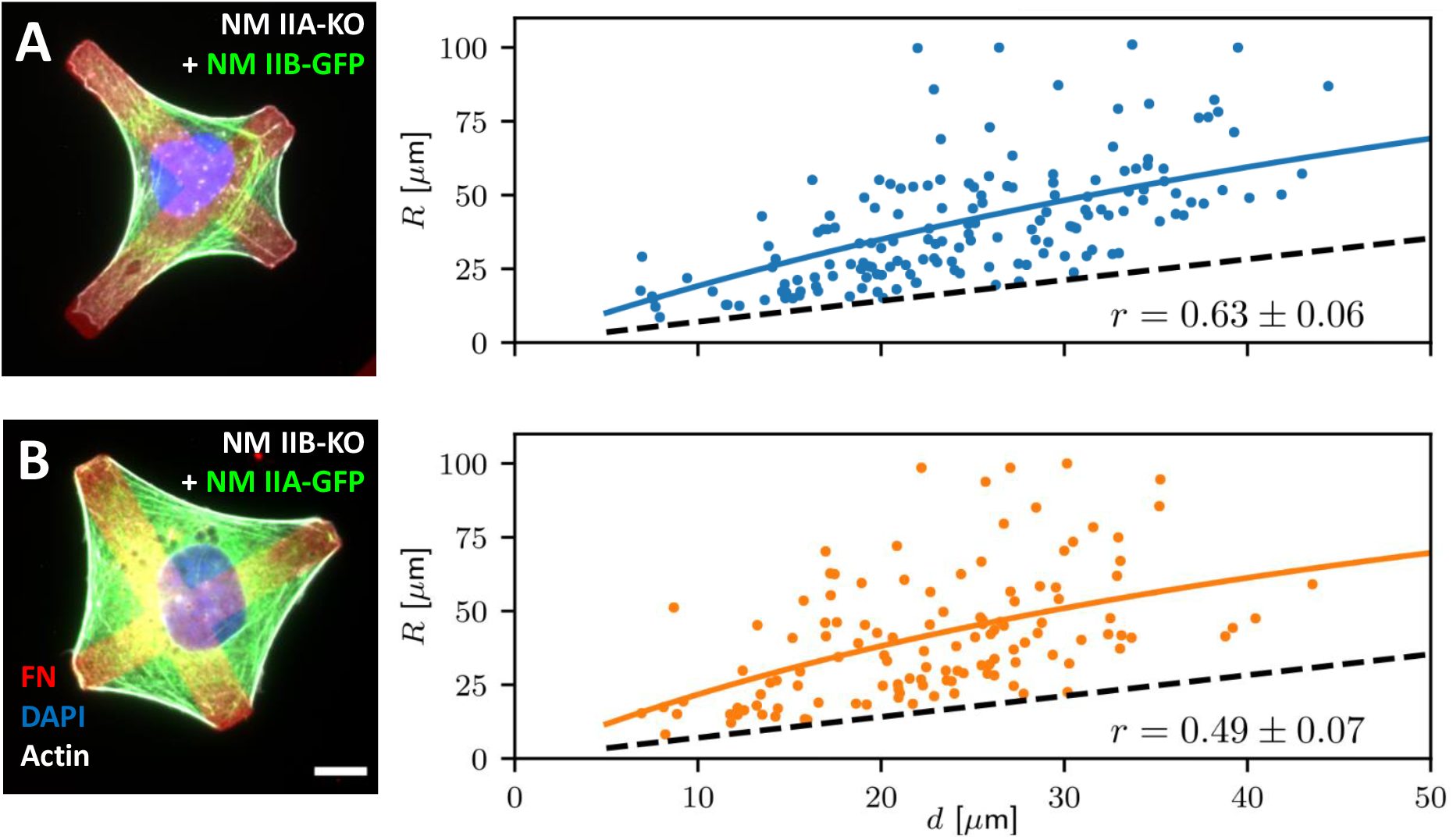
NM IIA or B overexpression on micropatterned substrates show the importance of the different motor qualities. **(A)** GFP-tagged NM IIB was overexpressed in NM IIA-KO cells and the cells were seeded on micropatterned substrates. Higher quantities of NM IIB did not restore the phenotype in the absence of NM IIA. The cells still spread along the cross bars rather than spanning large passivated areas and the *R*(*d*) ratio showed a positive correlation, comparable to WT and untransfected NM IIA-KO cells. **(B)** Overexpressing GFP-tagged NM IIA in NM IIB-KO cells did not restore the *R*(*d*) correlation, the values are comparably low to untransfected NM IIB-KO cells. Scale bars represent 10 µm for (A) and (B).

**Figure 4_figure supplement 1:**
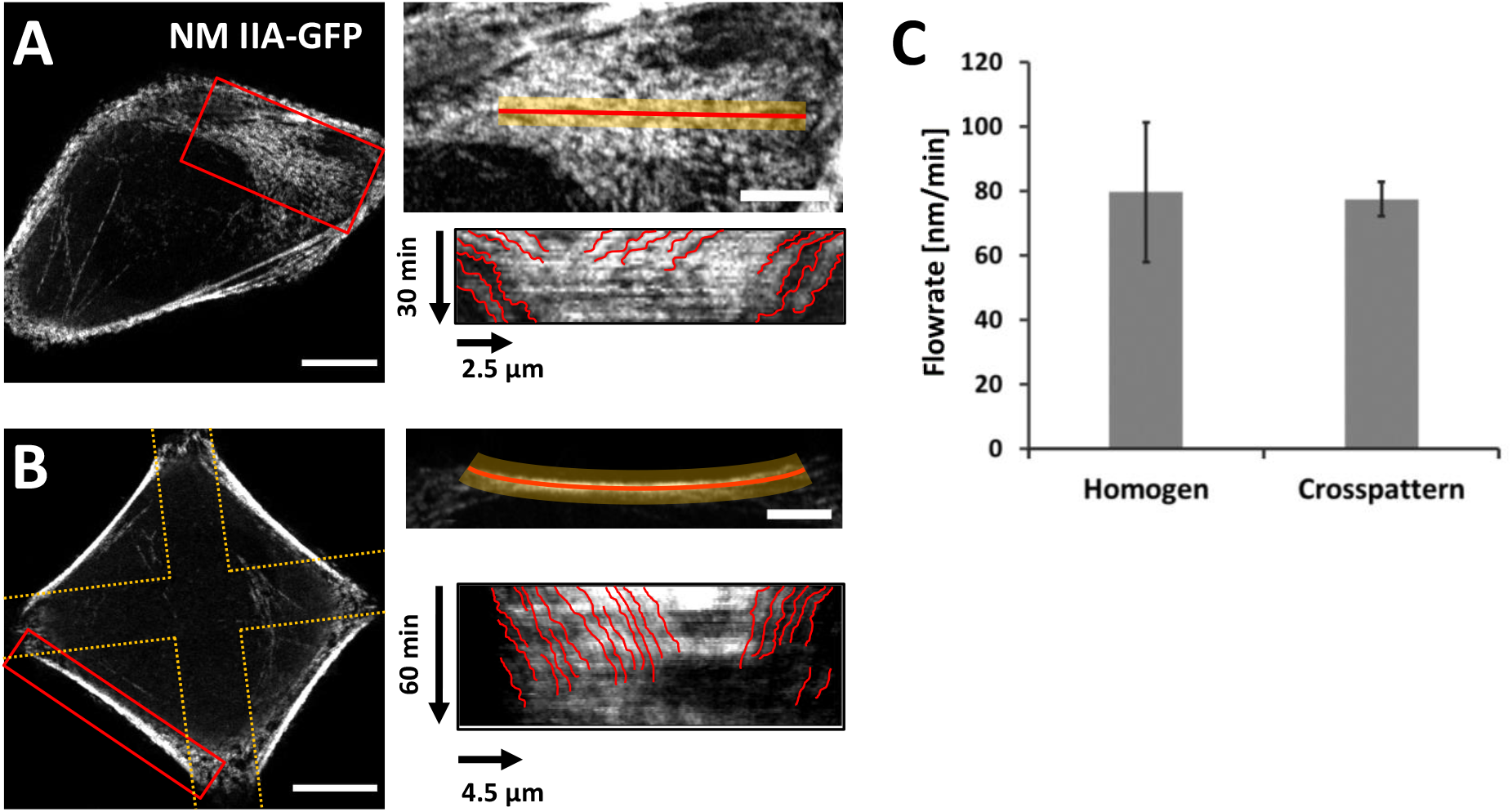
Cytoskeletal flow in actin SFs of cells on homogeneously coated substrates and in actin arcs of cells on cross-shaped micropatterns. U2OS WT cells were transfected with GFP-tagged NM IIA and cultivated on homogeneously coated FN-substrates **(A)** or FN-coated cross-shaped micropatterns **(B)**. Images were taken every minute for up to 1h using the AiryScan Mode. Kymographs were derived using the reslice mode of FIJI. The cytoskeletal flow in the kymograph was tracked manually and the flowrate was calculated from individual traces (depicted in red). **(C)** Mean values and standard deviations from all traces. Scale bars represent 10 µm in overviews and 5 µm in insets of (A) and (B).

**Figure 5_figure supplement 1:**
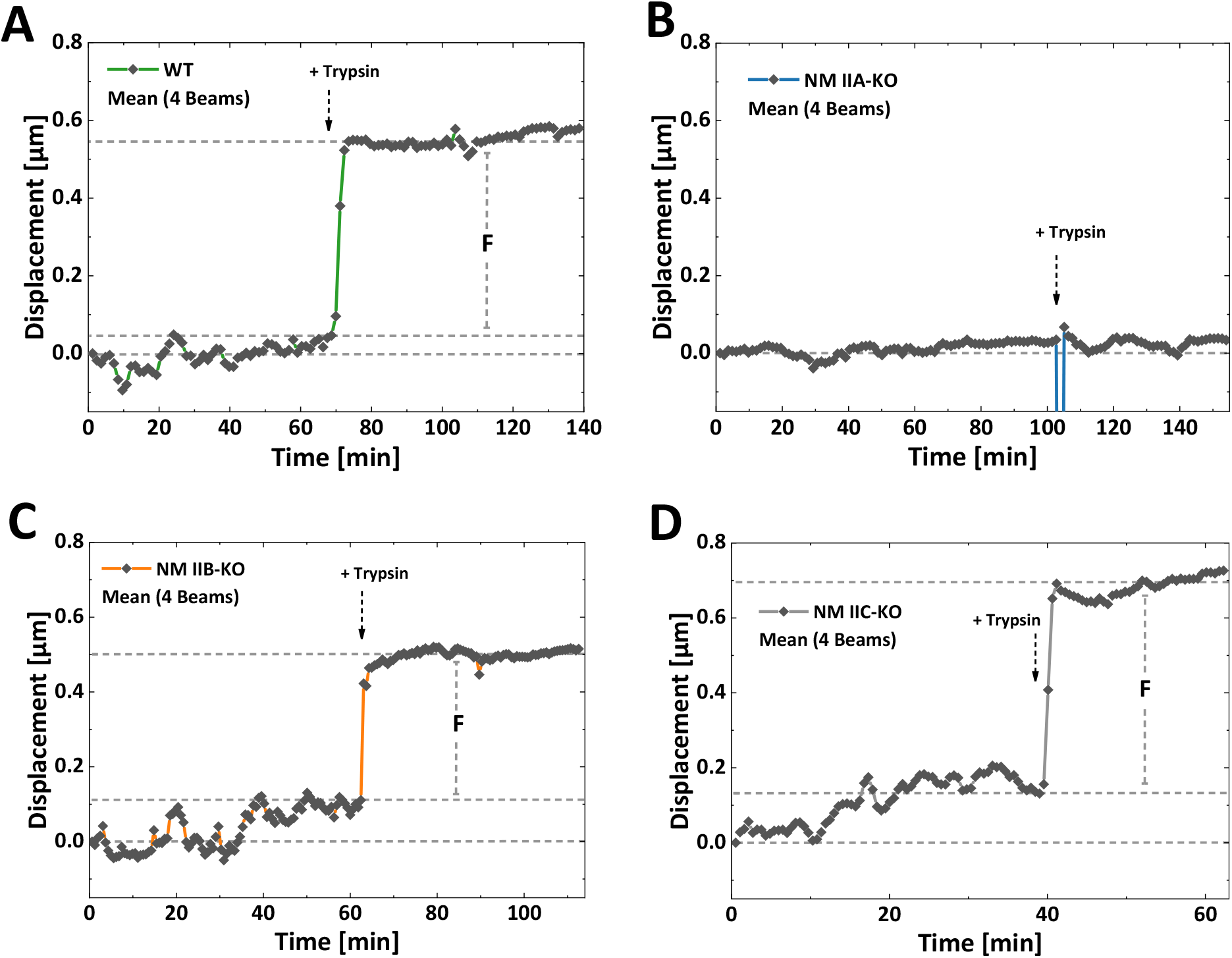
Initial force measurements of WT and NM II-KO cells. To measure initial forces, cells were seeded on 3D microscaffolds without stimuli-responsive hydrogels and the displacement of the adhesive bars was tracked before and after the addition of trypsin. In all graphs, dashed lines denote reference points at the start of the measurement, at the time point of trypsin addition and at the end of the experiment. Exemplary mean displacement traces of a single WT **(A)**, NM IIA-KO **(B)**, NM IIB-KO **(C)** and NM IIC-KO **(D)** cell correspond to Figure 5_supplement movies 1-4. No displacement was observed upon the addition of trypsin to NM IIA-KO cells, while WT, NM IIB-KO and NM IIC-KO cells each showed a striking displacement. Note that the displacement in (B) at ∼ 100 min resulted from focus issues in Figure 5_supplement movie 2.

**Figure 5_figure supplement 2:**
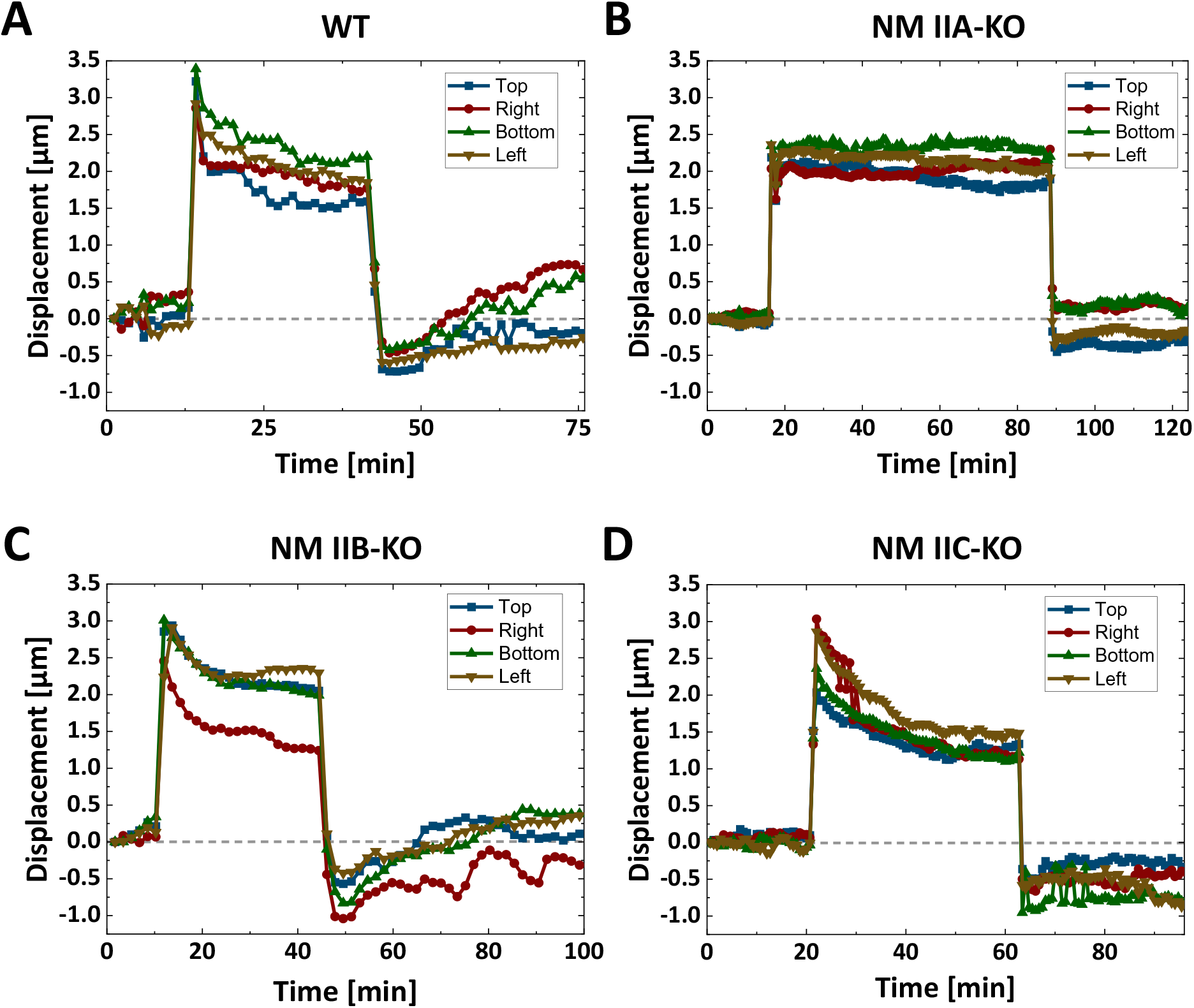
Displacements of individual beams for stretched WT and NM II-KO cells. Cells were seeded on stimuli-responsive 3D microscaffolds and the displacements of individual adhesive beams during the stretch-release cycle was tracked. **(A)** - **(D)** The displacement traces from this figure correspond to the mean displacement traces in figure 5A-D and figure 5_supplement movies 5-8. Dashed lines denote the reference point at the start of the measurement. **(A)** For the WT cell, all individual traces show a force increase after the stretch was applied and a force decrease after the stretch was released. **(B)** No force response was observed when stretching NM IIA-KO cells. **(C)** Individual beam displacements of a NM IIB-KO cell that show oscillatory motions in the cellular response, after the stretch was released (compare the red trace, corresponding to the right bar in figure 5_supplement movie 7). **(D)** Individual beam displacements of a NM IIC-KO, showing a force increase after the stretch but no force decrease during the time frame after the release.

