## supplemental text for "Distinct roles of nonmuscle myosin II isoforms for establishing tension and elasticity during cell morphodynamics"

### 1 Crossbridge cycle model for nonmuscle myosin II

In order to model the crossbridge cycle of the different isoforms of nonmuscle myosin II, we consider their three main mechanochemical states and the stochastic transitions between them, as depicted in Fig. S1A. Our three-state model has been extensively tested and parametrized before [1, 2, 3, 4] and is used here with a small modification. In brief, myosin heads bind from the unbound state to the actin filament with rate $k_{01} = 0.2\,\mathrm{s}^{-1}$. Recently it has been shown that this binding occurs in two steps, with a non-stereospecific state existing before the weakly bound state, which is stereospecific. The non-stereospecific intermediate state becomes relevant in the case of myosin II inhibition by blebbistatin [5], which we do not consider here, but this observation motivates us to choose a larger value $k_{10} = 0.4\,\mathrm{s}^{-1}$ for the unbinding rate from the weakly bound to the unbound state (rather than $0.004\,\mathrm{s}^{-1}$ as used before [1, 2, 3, 4]). From the weakly bound state, the powerstroke occurs with the high rate $k_{12} = 1.4 \cdot 10^6\,\mathrm{s}^{-1}$. Furthermore, the powerstroke is associated with swinging of the lever arm, which is simulated by increasing the individual motor strain $x_i$ by the powerstroke distance $d = 8\,\mathrm{nm}$. After having performed the powerstroke, myosin can either return to the weakly bound state with the relatively small rate $k_{21}$, or it can unbind from actin with a force- and isoform-dependent rate

$$k_{20}^{a/b}(F) = k_{20}^{0a/0b}[\Delta_c \exp(-k_m x_i/f_c) + (1-\Delta_c)\exp(k_m x_i/f_s)]. \tag{S1}$$

Here, $k_{20}^{0a/0b}$ are the transition rates at zero force of the A- and B-isoform respectively. In particular, the rate for isoform A $k_{20}^{0a} = 1.71\,\mathrm{s}^{-1}$ is much larger than the rate for isoform B $k_{20}^{0b} = 0.35\,\mathrm{s}^{-1}$, which constitutes the mechanochemical difference between isoforms A and B in our model. The two terms in the brackets represent two different unbinding pathways, namely the *catch-path* and the *slip-path*, respectively. Transitions along the catch-path become slower with increasing force with force scale $f_c = 1.66\,\mathrm{pN}$. The force on the motor is calculated by the motor stiffness $k_m$ and the motor strain $x_i$. Conversely, transitions along the slip-path become faster with increasing force with force scale $f_s = 10.35\,\mathrm{pN}$. $\Delta_c$ is the fraction of transitions following the catch-path at zero force, while the complement is the fraction of transitions following the slip-path. Together, these two pathways model a *catch-slip bond*, so dissociation decreases with force at low loading (*catch*) and increases again at high loading (*slip*). Force dependence of the rates is only taken into account if the motor is loaded against its direction of movement, the transition rate defaulting to $k_{20}^0$ for forces in the other direction. Note that in principle the force scales could also be modeled to depend on the isoform. *In-vitro* experiments suggest that the catch-path force scale for isoform B are lower than for isoform A [6]. Here they are kept equal for simplicity, as the difference in the rate at zero force is the dominating effect at low forces.

In a minifilament, the different motor heads are mechanically coupled to form a bipolar structures with two ensembles pulling in opposite directions. This tug-of-war situation is depicted in Fig. S1B. We assume that in each minifilament half, $N = 15$ motor heads are active [4]. They work against external springs with strains $z_-$ and $z_+$, respectively. In addition, each side of the minifilament consists of a variable number of NM IIA and NM IIB motors, $N_a^-$, $N_b^-$, $N_a^+$ and $N_b^+$, respectively, with $N_a^- + N_b^- = N$ and $N_a^+ + N_b^+ = N$. At all times the forces acting on the myosin heads of each sides are balanced against the forces in the external springs. This also yields the value for the motor strains $x_i$ required to evaluate Eq. S1.

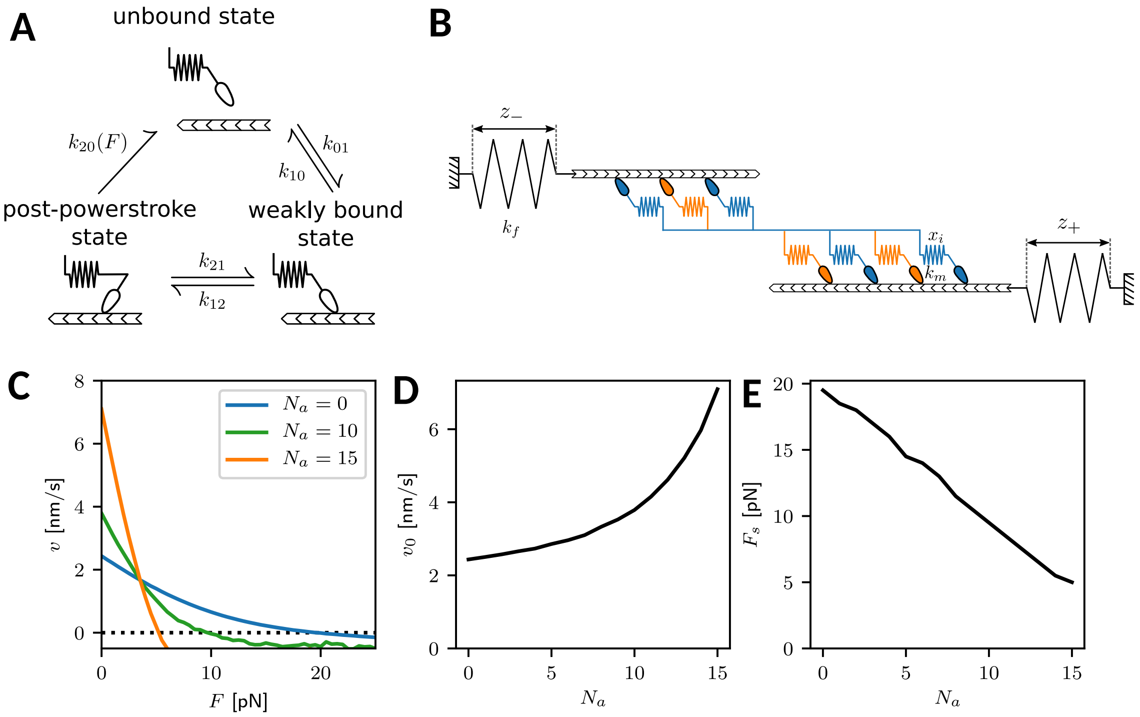


Figure S1: Crossbridge cycle model overview. (A) Mechanochemical crossbridge cycle for myosin II. (B) Tug-of-war of two mixed motor ensembles working against external springs. (C) Force-velocity relation for an ensemble (one half of a minifilament) with $N = 15$ motors with varying numbers $N_a$ of NM IIA motors. The number of NM IIB motors is $N_b = N - N_a$. (D) Force-free velocity $v_0$ and (E) stall force $F_s$ as a function of $N_a$.

All model parameters are summarized in Table S1. The stochastic model is simulated using the Gillespie algorithm, which uses the assumed rates to determine the time until the next reaction takes place [7]. From these simulations we obtain the force-velocity relations of the minifilaments with variable isoform content by averaging over many individual trajectories. As shown in Fig. S1C, we obtain well-defined relations in all cases, which allow us to predict the free velocity $v_0$ and the ensemble stall force $F_s$. We find that with increasing A-content, the free velocity $v_0$ increases and the stall force $F_s$ decreases, as shown in Fig. S1D and E, respectively. This demonstrates that minifilament with more A-isoforms are more dynamic, but also less stable mechanically. As a third important quantity, we can extract an effective friction coefficient $\xi_m$ from $v'(F_s) = -1/\xi_m$, which is the steepness of the force-velocity relation at the stall force. This friction coefficient $\xi_m$ decreases with increasing A-content, reflecting that the system becomes more dynamic.

### 2 Dynamic Tension Elasticity Model (dTEM)

The invaginated shapes of strongly adherent cells have been shown before to result from actomyosin contractility [8, 9, 10]. From a geometrical viewpoint, there are two actomyosin-related forces which balance each other along the invaginated arcs: an isotropic surface tension $\sigma$ in the cortex and a line tension $\lambda$ in the peripheral fiber. This leads to the Laplace law $R = \lambda/\sigma$ for the arc radius [9, 10]. Because experimentally it has been found that the radius $R$ also depends on the spanning distance $d$ of the invaginated arc, the line tension $\lambda$ has been argued to also contain an elastic contribution (tension-elasticity model, TEM), leading to $\lambda(d)$ and an increasing $R$-$d$ relation, as observed experimentally [10]. The line tension $\lambda$ can be identified with the force $F$ that acts in the fiber and in particular on the focal adhesions (FAs) by which it is anchored. Here we have confirmed the increasing $R$-$d$ relation and in addition revealed that it also depends on isoform content. In order to make contact to the different crossbridge cycles of isoform A versus B, we now introduce a dynamic variant of the TEM (dTEM).

We consider a peripheral fiber of length $L$ that is contracted by minifilaments that are distributed uniformely along its length. The contraction speed due to contractility is denoted by $v_c$

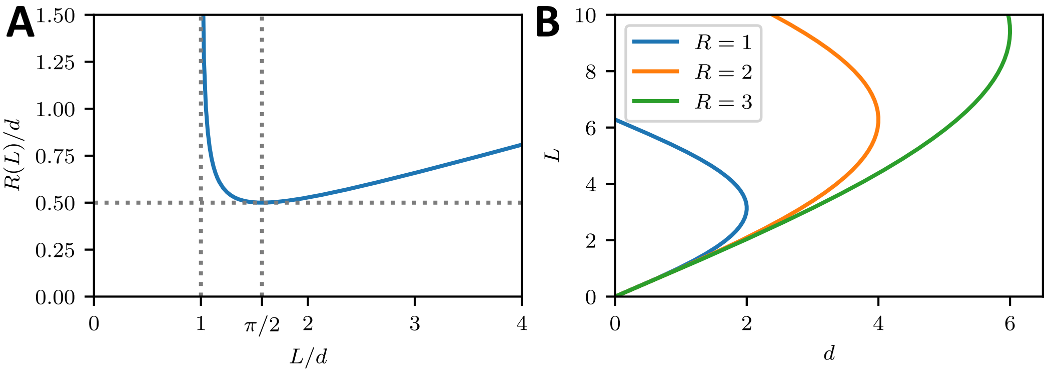


Figure S2: (A) The arc radius $R(L)$ is a function of arc length $L$ on the interval $L \in (d, \infty)$. For $L/d < \pi/2$ it is monotonically decreasing and for $L/d > \pi/2$ it is monotonically increasing. Normalizing $R$ and $L$ to the spanning distance $d$ yields a universal result. (B) Due to the geometry, $L(d)$ is not a well defined function, as for each spanning distance there exist two solutions for a central angle $0 < \varphi < 2\pi$.

and should depend on both the force $F$ inside the fiber and its length $L$ (like for muscle, a longer fiber should contract faster). In order to be able to obtain a stationary situation, new length has to be generated in the FAs at which the fiber is anchored, as observed experimentally as flow out of the FAs [11]. The corresponding polymerization velocity is named $v_p$ and also should depend on fiber force $F$, but not on $L$, because it is a local property of the FAs. Specifically, it has been shown that the main polymerization factor in FAs, the formin mDia1, increases its polymerization rate with force [12]. Together, the two velocities lead to a length change

$$\dot{L} = v_p(F) - v_c(F, L) \tag{S2}$$

of the stress fiber, which is the central dynamic equation that we will analyze here.

The contractile speed of molecular motors can be modeled by a linearized force-velocity relation

$$v_m(F) = \frac{1}{\xi_m}(F_s - F) \tag{S3}$$

with motor stall force $F_s$ and effective friction coefficient $\xi_m$ for a sarcomeric unit of reference length $L_0$ (for a linear force-velocity relation, two parameters are sufficient and the free velocity follows as $v_0 = F_s/\xi_m$). Linear scaling of contraction speed with stress fiber length implies for a stress fiber of length $L$

$$\frac{v_m}{L_0} = \frac{v_c(F, L)}{L} \;\Rightarrow\; v_c(F, L) = \frac{L}{L_0 \xi_m}(F_s - F). \tag{S4}$$

This relates the contraction speed in the fiber to the properties of the motor ensemble. For the polymerization speed of actin at the focal adhesions we assume a linear force dependence

$$v_p = \frac{F}{\xi_f} \tag{S5}$$

where $\xi_f$ is the effective friction coefficient inside the FAs.

The line tension that we obtain from the interplay of contraction and polymerization can now be related to the circular arc radius $R$ with the Laplace law

$$R = \frac{F}{\sigma} \ . \tag{S6}$$

This means that the dependence on $F$ in Eq. (S2) is in fact a dependence on the radius of curvature $R$, that in turn will depend on the length of the stress fiber $L$ and the spanning distance $d$. To close Eq. (S2), a geometrical equation that relates these quantities is required. Stress fiber length $L$ and the radius $R$ are trivially related by the central angle $\varphi$ as $L = R\varphi$. The central angle is in turn dependent on spanning distance $d$ and radius of curvature $R$. We then obtain

$$L = \begin{cases} 2R \arcsin\left(\frac{d}{2R}\right) & 0 \le \varphi \le \pi \\ 2R\left(\pi - \arcsin\left(\frac{d}{2R}\right)\right) & \pi \le \varphi \le 2\pi \end{cases} \tag{S7}$$

which is an implicit definition for $R(L,d)$. While circular arcs with larger central angle than $\pi$ cannot be observed on cross patterns, we do observe them on homogeneous substrates and in collagen gels. Therefore they are also included here in our discussion. In Fig. S2A we plot $R$ as a function of $L$ as defined by Eq. (S7) after rescaling all lengths with $d$. We see that the inversion is not unique and has two branches. This is due to $d(L)$ not being uniquely defined by Eq. (S7), which is visualized in Fig. S2B. For large $L/d$, the angle $\varphi$ is small, $R/d$ is large and $L \approx d$. For $\varphi = \pi$ (half-circle), we have $L/d = \pi/2$ and $R = d/2$. For larger $\varphi$, $R/d$ increases again and finally diverges. On the cross patterns, we typically deal with the small angle case.

By combining Eqs. (S2), (S4), (S5) and (S6), we now obtain our central dynamical equation:

$$\dot{L} = -\frac{L}{L_0 \xi_m}(F_s - \sigma R(L,d)) + \frac{\sigma R(L,d)}{\xi_f}. \tag{S8}$$

We non-dimensionalize this equation by measuring distance in units of $R_{\text{max}} = F_s/\sigma$ and time in units of $\tau = \xi_f/\sigma$:

$$\dot{l} = -\frac{l}{l_m}\left[1 - r(l,\delta)\right] + r(l,\delta) \tag{S9}$$

where $r$ is the dimensionless arc radius and $\delta$ is the dimensionless spanning distance. Moreover we have defined

$$l_m = \frac{\xi_m L_0}{\xi_f R_{\text{max}}} = \frac{\xi_m L_0 \sigma}{\xi_f F_s}\ . \tag{S10}$$

Fig. S3 shows $\dot{l}$ as a function of $l$. We see that for sufficiently small values of $\delta$ and $l_m$, both a stable and an unstable fixed point exist, at small and large values of $l$, respectively. The stable fixed point at small $l$ corresponds to a steady state. This state is lost through a saddle-node bifurcation for larger values of the dimensionless quantities $\delta$ and $l_m$.

By solving for the stationary state with $\dot{l} = 0$, we arrive at a relation between the arc radius $R$ and the steady state fiber length $L$, which we give both without and with dimensions:

$$r = \frac{l}{l_m + l}, \quad R = \frac{L}{L_m + L} R_{\text{max}}\ . \tag{S11}$$

Thus, the arc radius $R$ initially increases linearly with arc length $L$ and then starts to saturate towards $R_{\text{max}} = F_s/\sigma$ at the arc length of half maximal radius $L_m = \xi_m L_0/\xi_f$. Here, the two competing length scales are the Laplace radius $R_{\text{max}}$ and the ratio of the slopes of the length-normalized force-velocity relation of the fiber and the force-velocity relation of the FA.

To compare with our experimental results, we finally have to convert the $R(L)$ relation to a $R(d)$ relation. Practically this can be done best by first fixing $d$, then finding the lower fixed point for $l$ of Eq. S9 (if it exists) and finally calculating $R$ from Eq. S11. Alternatively, one can use the approximation $L \approx d$ for small central angles, which we did in the main text. We then arrive at

$$r = \frac{\delta}{\delta_m + d}, \quad R = \frac{d}{d_m + d} R_{\text{max}}, \tag{S12}$$

where we have renamed $l_m = \delta_m$ and $L_m = d_m$. This equation is the central result (1) in the main text. The result for $\delta_m < \pi$ is shown in Fig. S4A. For the exact case we find $R(d)$ relations that always start at $R(0) = 0$. From there, the function increases monotonically while curving downward and approaching a plateau for low $\delta_m$. In this regime, the approximation is reasonably accurate. At $d_{\text{high}}$, where $\varphi = \pi$, the function stops approaching the plateau. Shortly after, at $d_{\text{crit}}$, the saddle-node bifurcation occurs and at higher spanning distances $d$ no steady state exists anymore. For the parameters chosen in Fig. S4A, $d_{\text{high}} \approx d_{\text{crit}}$ and for this reason, the region where central angles $\varphi > \pi$ is very small.

For $\delta_m \ge \pi$, the central angle $\varphi > \pi$. Again this cannot be observed on cross patterns, but is discussed here for completeness. The $R(d)$ relation again starts at zero for zero spanning

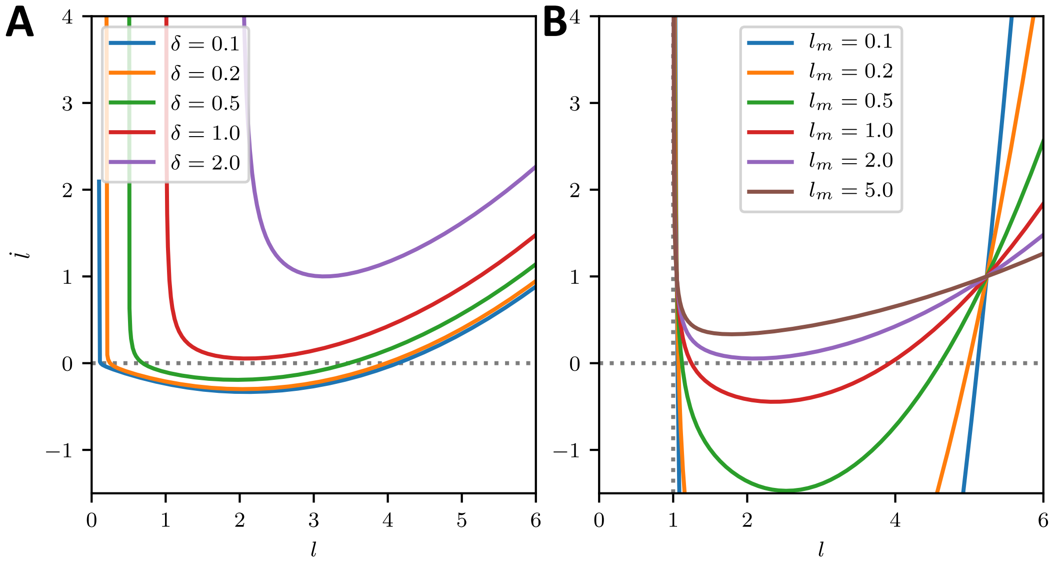


Figure S3: Stability analysis. (A) Change in dimensionless arc length $\dot{l}$ for a range of spanning distances $\delta$ as a function of $l$ for $l_m = 2$. For small spanning distances there are two steady states. The one at lower $l$ is stable, while the other one is unstable. For increasing $\delta$ the system approaches a saddle-node distribution, beyond which both steady states vanish and the length of the peripheral arc increases indefinitely. (B) $\dot{l}$ for a range of $l_m$ and spanning distance $\delta = 1$. Again the two fixed points vanish in a saddle node bifurcation.

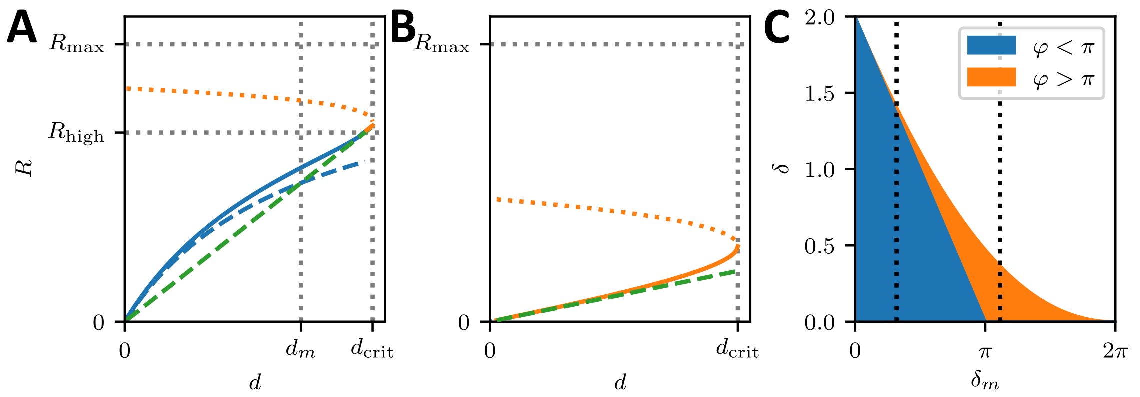


Figure S4: (A,B) $R(d)$ relationship predicted by the model. Blue solid lines are related to stable steady state lengths, where the arc is smaller than a half cicle, while dashed blue lines represent the result with approximation $\arcsin x = x$. Solid orange lines represent stable steady states where the central angle $\varphi > \pi$ and dotted orange lines represent unstable steady states. The dashed green line denotes $d = 2R$, which corresponds to the circle of smallest radius by geometry. The colors in the phase diagrams represent parameter regions, where stable solutions can be found and represent arcs that are smaller (blue) and larger (orange) than half circles, respectively. In the white area the arc curls inward and increases its radius indefinitely. (A) $\delta_m = 1$ (B) $\delta_m = 3.5$ (C) Phase diagram indicating whether the central angle is larger or smaller than $\pi$. The dashed vertical lines visualize the parameter range described by subfigure A and B respectively.

distance and curves upward until reaching the critical value for $d$ where the steady states do not exist anymore as shown in Fig. S4B. Fig. S4C gives a general overview of the parameter range indicating regions of upward curvature connected to central angles lower than $\pi$ together with regions of downward curvature which are connected to central angles higher than $\pi$.

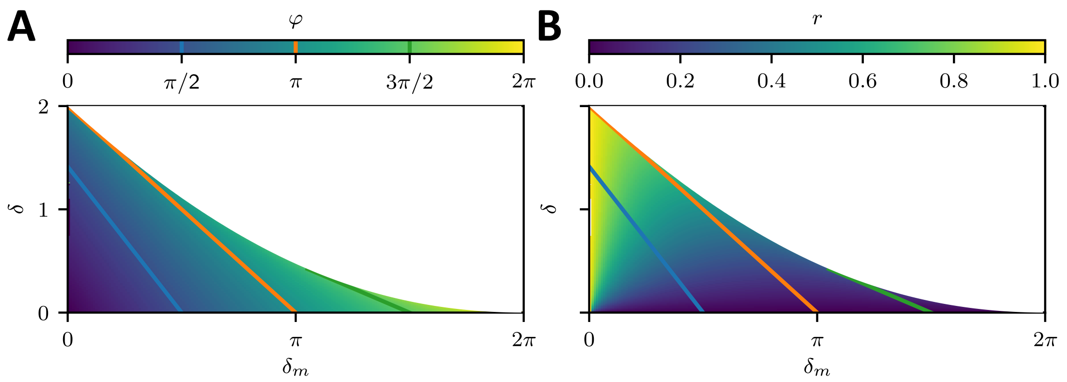


Figure S5: Phase diagram. (a) Central angle $\varphi$ and (b) dimensionless radius $r$ as a function of $\delta$ and $\delta_m$. The white region represents unstable parameter configurations, while the colored areas represent the stable steady state central angle and radius respectively. The colored lines are contour lines of the central angle and indicate upper bounds for stability given that the maximum permissible central angle by the micropattern geometry.

Using $\varphi r = l$ and $\sin\varphi/2 = \delta/2r$ in Eq. (S11) we find

$$\delta = 2\sin\frac{\varphi}{2}\left(1 - \frac{\delta_m}{\varphi}\right), \tag{S13}$$

i.e. the contour lines of the central angle $\varphi$ as a function of $(\delta, \delta_m)$ are linear functions. In particular for $\varphi = \pi$, which is the situation of highest spanning distance given the radius, we find:

$$d_{\text{high}} = 2(R_{\max} - \frac{d_m}{\pi}). \tag{S14}$$

For angles $\varphi > \pi$ the linear functions from eq. (S13) however intersect, which contour lines should never do. This implies, that the contour lines for $\varphi$ have to be constrained more carefully. As the steady state central angle $\varphi(\delta_m, \delta)$ is a smooth function that is well defined for low enough $\delta_m$ and $\delta$ and monotonically increases with $\delta_m$ and $\delta$ we can search for the maximum stable spanning distance $\delta$ w.r.t. central angle $\varphi$ using eq. (S13). This yields,

$$\begin{aligned}\delta_m^{\max} &= \frac{\varphi^2\cos\frac{\varphi}{2}}{\varphi\cos\frac{\varphi}{2} - 2\sin\frac{\varphi}{2}} \\ \delta^{\max} &= \frac{\left(2\sin\frac{\varphi}{2}\right)^2}{2\sin\frac{\varphi}{2} - \varphi\cos\frac{\varphi}{2}}\end{aligned}, \tag{S15}$$

which marks the border of stability of the system for $\pi \leq \varphi \leq 2\pi$ as shown in Fig. S5. Contour lines of $\varphi$ for $\varphi > \pi$ start on this curve, with smaller $\delta_m$ having to be excluded from Eq. (S13).

Fig. S5 shows the overall dependencies of the central angle $\varphi$ and the radius $r$ on the spanning distance and the respective length scale $\delta_m$ together with contour lines for the central angle $\varphi$, which can also be interpreted as upper bounds for stability given a specific micropattern geometry. The cross shaped micropattern yields maximum central angles of $\pi/2$ (blue line in Fig. S5 and upper bound of the red region in Fig. 3E of the main text) as illustrated in Fig. 3B of the main text.

Table S1: Model parameters.

| Parameter | Symbol | Value | References |
|---|---|---|---|
| Transition rates [$s^{-1}$] | $k_{20}^{a0}$ | 1.71 | [3, 13] |
| | $k_{20}^{b0}$ | 0.35 | [3, 13] |
| | $k_{01}$ | 0.2 | [3, 13] |
| | $k_{10}$ | 0.4 | [5] |
| | $k_{12}$ | $4 \cdot 10^6$ | [3, 14] |
| | $k_{21}$ | 0.7 | [3, 14] |
| Catch-path fraction | $\Delta_c$ | 0.92 | [3, 13] |
| Catch-path force scale [pN] | $f_c$ | 1.66 | [3, 13] |
| Slip-path force scale [pN] | $f_s$ | 10.35 | [3, 13] |
| Neck-linker stiffness [pN/nm] | $k_m$ | 0.7 | [3, 13] |
| Powerstroke distance [nm] | $d$ | 8 | [1, 2, 14] |
| External springs [pN/nm] | $k_f$ | 4 | [15] |
