## Supplementary figures and images for "Distinct roles of nonmuscle myosin II isoforms for establishing tension and elasticity during cell morphodynamics"

### Figure 4_movie 1

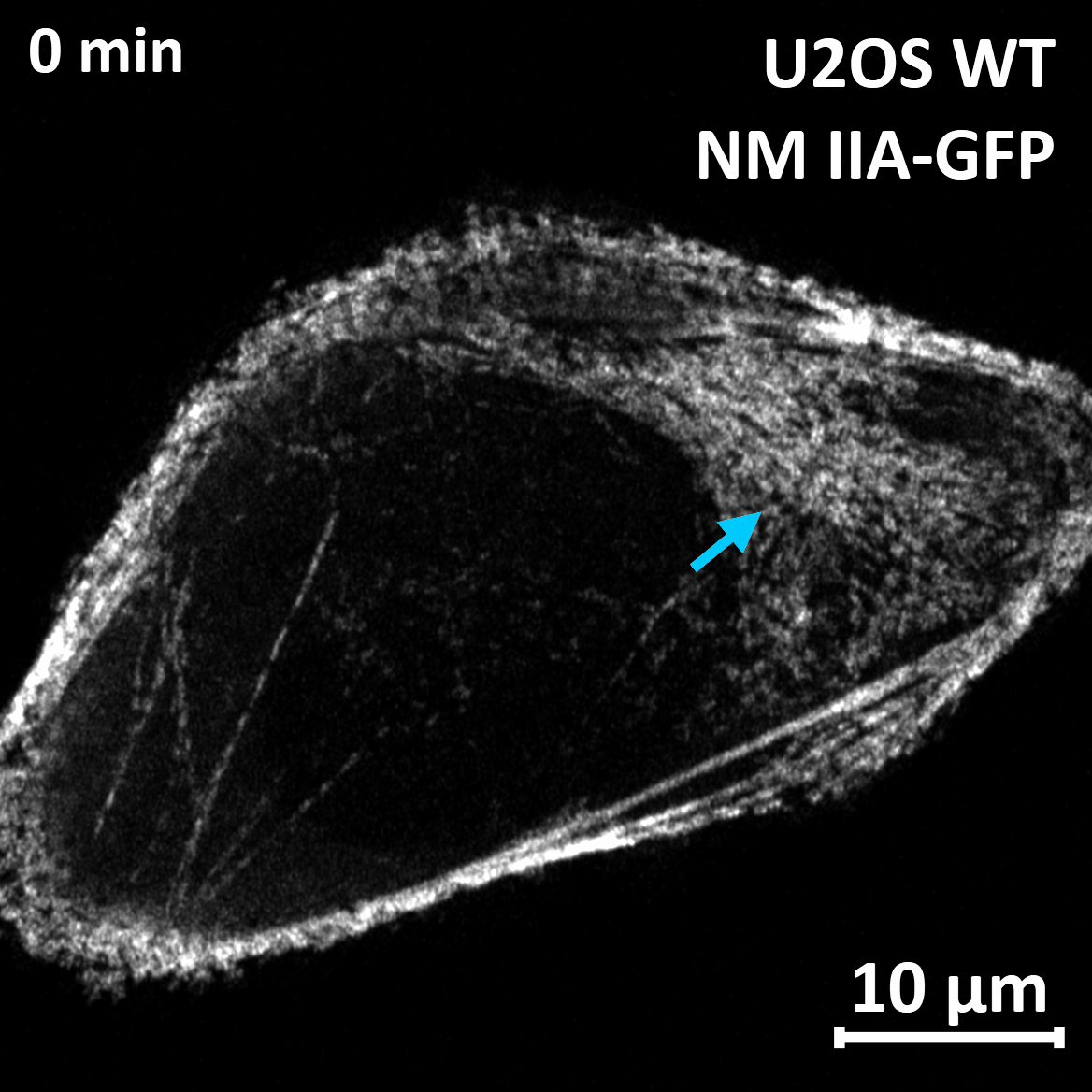

### Figure 4_movie 2

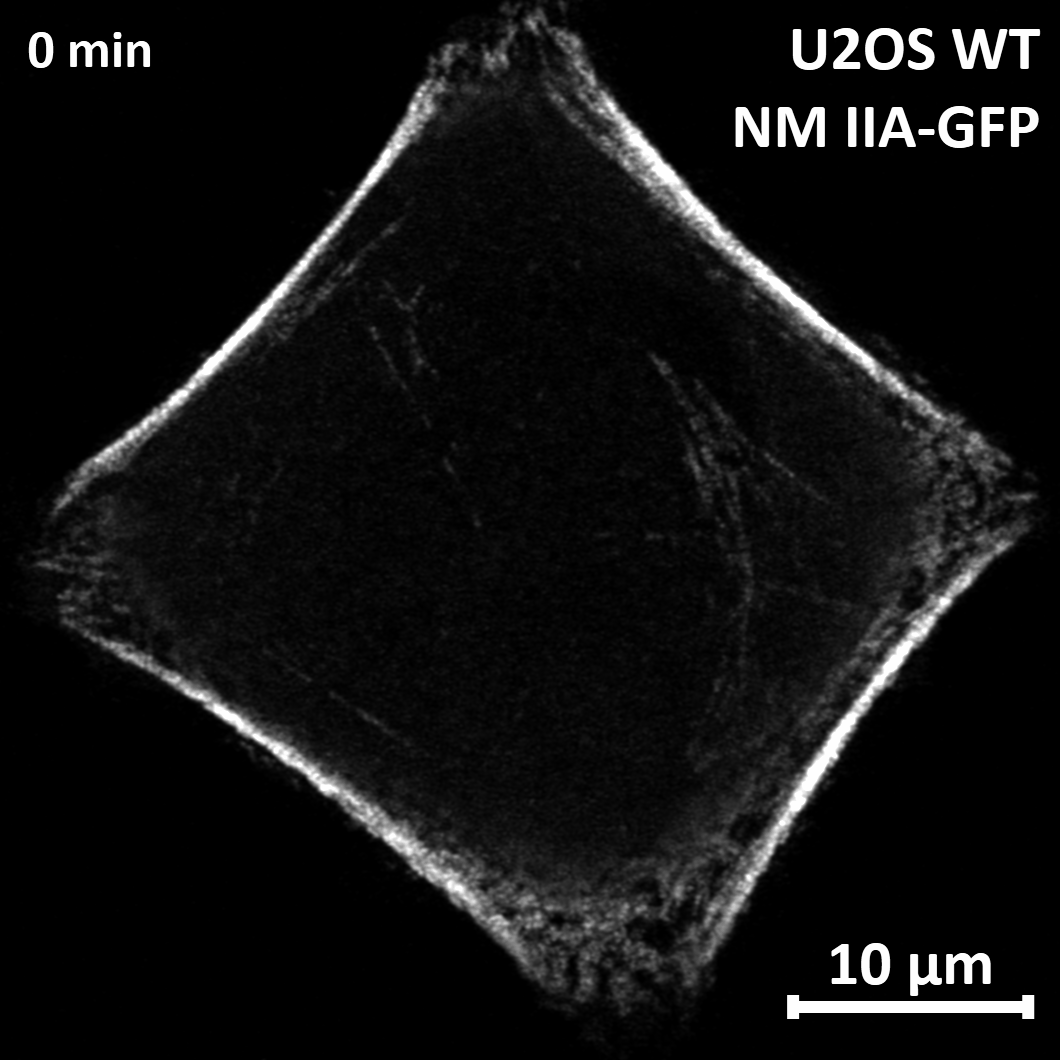
